# Vimentin Intermediate Filaments Stabilize Dynamic Microtubules by Direct Interactions

**DOI:** 10.1101/2020.05.20.106179

**Authors:** Laura Schaedel, Charlotta Lorenz, Anna V. Schepers, Stefan Klumpp, Sarah Köster

## Abstract

The cytoskeleton determines cell mechanics and lies at the heart of important cellular functions. Growing evidence suggests that the manifold tasks of the cytoskeleton rely on the interactions between its filamentous components – actin filaments, intermediate filaments and microtubules. However, the nature of these interactions and their impact on cytoskeletal dynamics are largely unknown. Here, we show in a reconstituted in vitro system that vimentin intermediate filaments stabilize microtubules against depolymerization and support microtubule rescue. To understand these stabilizing effects, we directly measured the interaction forces between individual microtubules and vimentin filaments. Combined with numerical simulations, our observations provide detailed insight into the physical nature of the interactions and how they affect microtubule dynamics. Thus, we describe an additional, direct mechanism for cells to establish the fundamental cross-talk of cytoskeletal components alongside linker proteins. Moreover, we suggest a novel strategy to estimate the binding energy of tubulin dimers within the microtubule lattice.

---

The cytoskeleton is a dynamic biopolymer scaffold present in all eukaryotic cells. Its manifold tasks depend on the fine-tuned interplay between the three filamentous components, actin filaments, microtubules and intermediate filaments (IFs).^1–6^For example, all three types of cytoskeletal polymers participate in cell migration, adhesion and division.^3–6^ In particular, the interplay of IFs and microtubules makes an important contribution to cytoskeletal cross-talk, although the interaction mechanisms largely remain unclear. ^1,7–18^

For instance, vimentin, one of the most abundant members of the IF family, forms closely associated parallel arrays with microtubules in migrating cells.^7,16,18^Depolymerization of the microtubule network leads to a collapse of vimentin IFs to the perinuclear region, further attesting their interdependent organization in cells.^9^ Several studies suggest that in cells, microtubules associated with the vimentin IF network are particularly stable: They exhibit increased resistance to drug-induced disassembly ^9^ and enhanced directional persistence during directed cell migration,^18^and they are reinforced against lateral fluctuations.^17^ Several proteins such as kinesin, ^8,11^dynein,^13,15^plectin^1^and microtubule-actin cross-linking factor (MACF)^10,12^can mediate interactions between IFs and microtubules. These linker proteins may be involved in conferring microtubule stability in cells. However, the possibility that more fundamental, direct interactions independent of additional components like microtubule associated proteins may contribute to stabilizing microtubules remains unexplored. Such a mechanism could also explain the results of an in vitro study on dynamic microtubules embedded in actin networks: Depending on the network architecture, actin regulates micro-tubule dynamics and life time. In particular, unbranched actin filaments seem to prevent microtubule catastrophe, thus stabilizing them, though the exact interaction mechanism is not revealed.^19^ In contrast to the cell experiments that showed a stabilization of microtubules by IFs, earlier work found that many IFs, including vimentin, contain tubulin binding sites and that short peptides containing these binding sites inhibit microtubule polymerization in vitro.^14^ Yet, it is unknown how this effect relates to fully assembled vimentin filaments.

Overall, there is substantial evidence for the importance of the interplay between IFs and microtubules in cells. However, the nature of their direct interactions and its impact on cytoskeletal dynamics remains elusive. Here, we studied these interactions by combining in vitro observations of dynamic microtubules in presence of vimentin IFs with single filament interaction measurements and complementary numerical simulations. In stark contrast to Ref. 14, our observations and simulations of dynamic microtubules reveal a stabilizing effect by the surrounding vimentin IFs. Based on our experimental data, we also estimated the tubulin dimer binding energy within the microtubule lattice, which is a much sought-after parameter for understanding microtubule dynamic instability. ^20–27^ This value has previously only been determined by molecular dynamics simulations and kinetic modeling^20,22,27^ or by using atomic force microscopy to indent stabilized microtubules.^24^

To study the influence of IFs on microtubule dynamics, we polymerized microtubules in the presence of vimentin IFs. We imaged the microtubules by total internal reflection fluorescence (TIRF) microscopy as sketched in Fig. 1a. As nucleation sites for dynamic microtubules, we used GMPCPP-stabilized microtubule seeds (green in Fig. 1a) adhered to a passivated glass surface. For simultaneous assembly of microtubules (cyan) and IFs (red), we supplemented a combined buffer (CB) containing all ingredients necessary for the assembly of both filament types with 20 µM or 25 µM tubulin dimers and 2.3 µM or 3.6 µM vimentin tetramers (0.5 or 0.8 g/L protein). All experiments presented in this work refer to these protein concentrations. All TIRF experiments were carried out in the same buffer conditions. Fig. 1b shows a typical fluorescence image of mixed microtubules and vimentin IFs.

**Figure 1:**
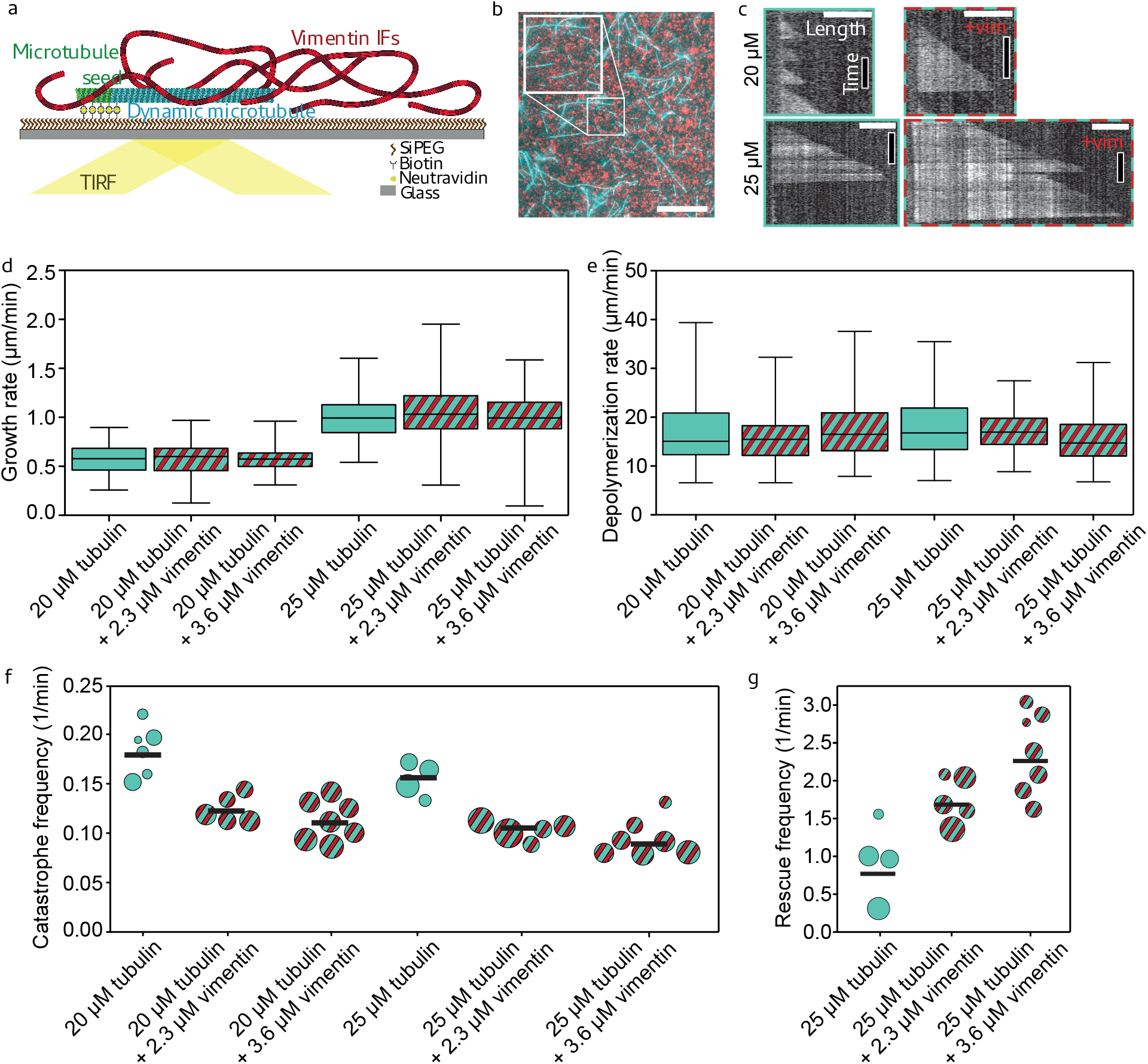
Vimentin IFs stabilize dynamic microtubules. (a) Illustration of the experimental setup. We attached microtubule seeds (green) to a biotin-SiPEG coated cover glass. Dynamic microtubules (cyan) grew from the seeds. Vimentin IFs (red) formed an entangled, fluctuating network. We imaged microtubules by TIRF microscopy. (b) Fluorescence micrograph of microtubules (cyan) embedded in a vimentin IF network (red). Inset shows enlarged detail. Scale bar: 10 µm. (c) Example kymographs of microtubules growing at 20 µM or 25 µM tubulin in the presence (+vim; 2.3 µM) or absence of vimentin. Scale bars: 3 µm and 5 min. (d) and (e) Vimentin does not affect the microtubule growth and depolymerization rates, irrespective of the tubulin concentration. Cyan boxplots represent experiments with tubulin only, cyan and red striped boxplots illustrate experiments with tubulin and vimentin. Boxplots include the median as the center line, the 25th and 75th percentiles as box limits and the entire data range as whiskers. (f) The catastrophe frequency of microtubules decreases in the presence of vimentin. Each circle represents an experiment including multiple microtubules. The circle area scales with the total summed microtubule growth time of the respective experiment. Black bars indicate the weighted mean. (g) Vimentin enhances the microtubule rescue frequency. Each circle represents an experiment including multiple microtubules. The circle area scales with the total microtubule depolymerization time. All tubulin and vimentin concentrations are input concentrations.

We analyzed microtubule dynamics using kymographs obtained from TIRF microscopy as shown in Fig 1c. As expected,^28^ the microtubule growth rate increased at the higher tubulin concentration (Fig. 1d, cyan). Yet, the presence of vimentin IFs did not affect the growth and depolymerization rates (Fig. 1d, e). Interestingly, we observed a marked decrease in the catastrophe frequency^29^ in presence of vimentin IFs at both tubulin concentrations (Fig. 1f, red and cyan stripes). Moreover, vimentin IFs promote microtubule rescue (Fig. 1g). As rescue events are rare at the lower tubulin concentration, ^29^ we only report rescue data for 25 µM tubulin. When assembly is initiated, vimentin unit-length filaments form after about 100 ms.^30^Therefore, we assume that vimentin filaments, not precursors, interacted with the microtubules. Additionally, we did not observe differences in microtubule dynamics when comparing early and late time points within the same experiment, although the mean vimentin filament length increased over the course of the experiments (see supplementary Fig. 1). These results indicate that vimentin IFs stabilize dynamic microtubules by suppressing catastrophe and enhancing rescue, while leaving the growth rate unaffected. A higher vimentin concentration enhances these effects.

From these observations, we hypothesize that there are direct, attractive interactions between microtubules and vimentin IFs, which stabilize dynamic microtubules. To test this hypothesis, we studied the interactions of single stabilized microtubules and vimentin IFs with optical trapping (OT), a complementary method to our TIRF experiments, as illustrated in Fig. 2. We prepared fluorescent and biotin-labeled microtubules and vimentin IFs as sketched in Supplementary Fig. 2. We used an OT setup, combined with a microfluidic device and a confocal microscope (LUMICKS, Amsterdam, The Netherlands) to attach a microtubule and a vimentin IF to separate bead pairs via biotin-streptavidin bonds as shown in Fig. 2a and Supplementary Fig. 3a. Once the IF and microtubule were in contact, we moved the IF back and forth in *y*-direction. If the IF and microtubule interacted, eventually either the IF-microtubule interaction broke (Fig. 2b) or the IF-microtubule interaction was so strong that the microtubule broke off a bead (Fig. 2c). To study the orientation dependence of the interaction, we included two additional measurement geometries: (i) we turned the microtubule by 45° as shown in Fig. 2e or (ii) moved the IF horizontally in the *x*-direction along the microtubule (Fig. 2f). We categorized the type of interaction, i.e. no interaction, IF-microtubule bond broke, or microtubule broke off bead, for each filament pair, as shown by pictograms in Fig. 3a, top.

**Figure 2:**
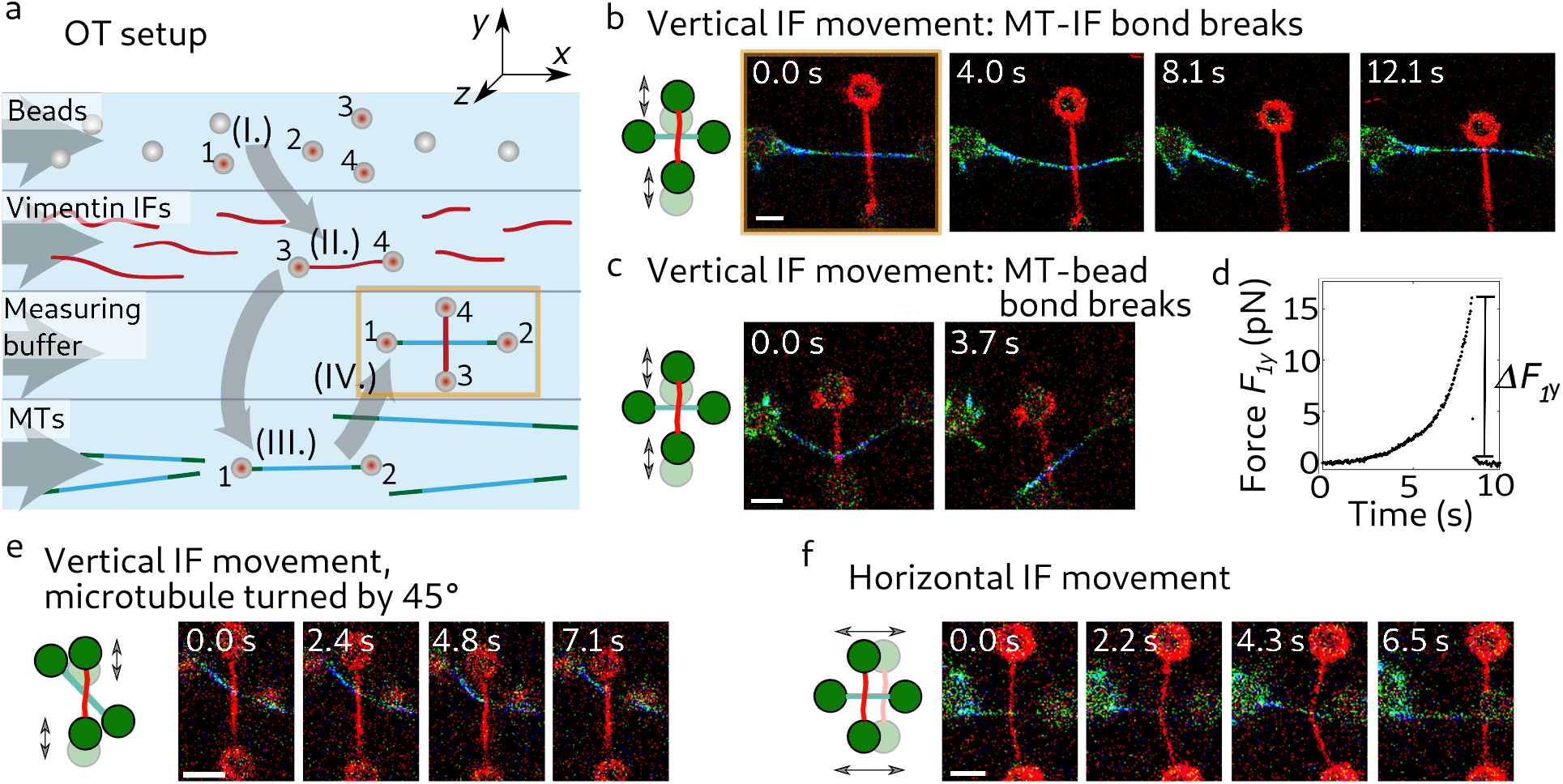
Direct interactions between stabilized microtubules and vimentin IFs. (a) Schematic of the setup for the OT experiments for interaction measurements in microfluidic flow channels. Four streptavidin-coated beads were captured by OTs (I.). We used one bead pair (beads 3 and 4) to attach a vimentin IF (II., red), and the other bead pair (beads 1 and 2) to attach a microtubule (III., green-cyan). We brought the IF and the microtubule into contact in a crossed configuration (IV.). Next, we moved the IF perpendicularly to the microtubule to study the IF-microtubule interactions while we took confocal fluorescence images (starting position marked in yellow in (a) and (b)). (b) Typical confocal fluorescence images of an IF-microtubule interaction which broke while the IF was moved vertically and (c) a strong IF-microtubule interaction for which the microtubule broke off the bead. (d) Typical experimental force increase *F*_1*y*_ on bead 1 in the *y*-direction once a bond forms. Breaking of the force causes a force jump of ∆*F*_1*y*_. (e) Typical confocal fluorescence images of a breaking IF-microtubule interaction at a 45°angle between them while the IF was moved vertically. (f) Typical confocal fluorescence images of a breaking IF-microtubule interaction in perpendicular configuration while the IF was moved horizontally. Scale bars: 5 µm.

**Figure 3:**
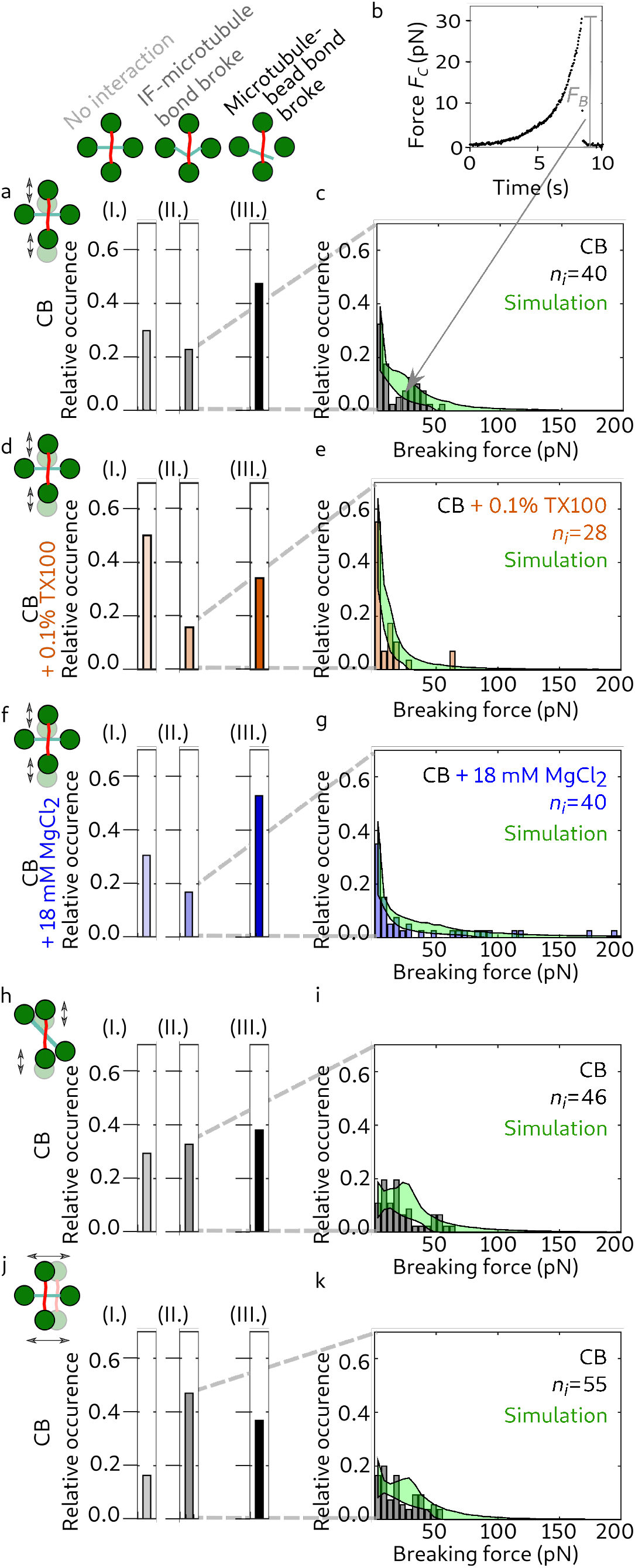
Hydrophobicity and electrostatics contribute to the IF-microtubule interactions. We classified the interactions between IF-microtubule pairs into three different groups as shown by the pictograms: no interaction (I.), breaking of the IF-microtubule bond as shown in Fig. 2b, e and f (II.), and breaking of the microtubule-bead bond as shown in Fig. 2c (III.). Typical experimental force-time behavior of the IF-microtubule bond showing the total force acting on the IF-microtubule bond, *F*_*C*_. The plot represents the corrected version of the force data shown in Fig. 2d taking into account the geometry of the filament configuration. Histograms of *n*_*i*_ experimentally recorded breaking forces (gray) and simulated data (green) for measurements in pure CB when the IF-microtubule interaction broke as shown in Fig. 2b. Due to statistical fluctuations, the distribution appears to be bimodal, however it can still be well described with a unimodal distribution. (d) TX100 (orange) suppresses some of the interactions, which results in more IF-microtubule pairs without any interaction (aI. vs. dI.) and fewer instances of IF-microtubule interactions (aII.-III. vs. dII.-III.). (e) The IF-microtubule bonds formed in presence of TX100 break at lower forces. (f) Magnesium (blue) does not change the relative number of IF-microtubule pairs which do not interact (aI. vs. fI.), but leads to fewer IF-microtubule breaking events (aII. vs. fII.) because the interactions become so strong that the microtubule breaks off the bead more often (aIII. vs. fIII.). (g) The IF-microtubule bonds formed in presence of additional magnesium break at higher forces. (h) When the microtubule was turned by 45°, IF and microtubule interacted more frequently (hII vs. aII). (i) The corresponding distribution of the breaking forces resembles the distribution for the perpendicular geometry (see c). (j, k) The binding rate for a horizontal movement of the IF increases compared to a vertical movement (jII vs. aII). (k) The corresponding distribution of breaking forces is similar to the distributions for the other two geometries (see i and c).

With the OTs, we recorded the force *F*_1*y*_ or *F*_1*x*_ that acted on trap 1 (see Fig. 2d and Supplementary Fig. 3b), which increased after the IF bound to the microtubule. Based on the geometry of the filament configuration from the confocal images, we calculated the total force *F*_*C*_ that the IF exerted on the microtubule (see Supplementary Fig. 3c). In Fig. 3b we show the resulting force calculated for the data shown in Fig. 2d. Combining all experiments with a breaking IF-mcirotubule bond leads to a distribution of *n*_*i*_ breaking forces *F*_*B*_ as shown in the force histogram in Fig. 3c. Due to thermal fluctuations, the force resolution of our system is limited to 1 pN and we thus focused on interaction forces above 1 pN, which is consistent with physiologically occurring intracellular forces. The breaking forces are in the range of 1-65 pN with higher forces occurring less often. Hence, in agreement with our hypothesis, our experiments show that single microtubules and vimentin IFs directly interact, i.e. without involving any linker proteins, and that these interactions can become so strong that forces up to 65 pN are needed to break the bonds. This range of forces is physiologically relevant and comparable to other microtubule associated processes: Single microtubules can generate pushing forces of 3-4 pN while forces associated with depolymerization can reach 30-65 pN.^31^ Kinesin motors have stalling forces on the order of a few pN.^32^

To better understand the nature of the interactions between single microtubules and vimentin IFs, we varied the buffer conditions in which we measured filament interactions. First, we probed possible hydrophobic contributions to the interactions by adding 0.1% (w/v) Triton-X 100 (TX100), a non-ionic detergent. Rheological studies of IF networks previously suggested that TX100 inhibits hydrophobic interactions. ^33^ Tubulin dimers have several hydrophobic regions as well, ^34^ some of which are accessible in the assembled state.^35^ As shown in Fig. 3d and e, the number of interactions decreases and the breaking forces are slightly lower in presence of TX100 than in pure CB. We calculated the binding rate *r*_*b*,eff_ by dividing the total number of interactions larger than 1 pN by the time for which the two filaments were unbound: TX100 leads to a lower binding rate *r*_*b*,eff,TX100_ = 0.56 · 10^−2^ s^−1^ compared to the binding rate *r*_*b*,eff,*y*_ = 1.1 · 10^−2^ s^−1^without TX100. We speculate that TX100 interferes with the binding sites on both filament types by occupying hydrophobic residues on the surface of the filaments and thereby inhibits hydrophobic interactions between the biopolymers.^33^Consequently, the reduced number of interactions in presence of TX100 indicates that hydrophobic effects contribute to the interactions.

Next, we tested for electrostatic contributions to the interactions by adding magnesium chloride to the buffer. When probing interactions in CB buffer with a total concentration of 20 mM magnesium, we observed both an increase in strong interactions, where the IF pulls the microtubule off a bead, and higher breaking forces (Fig. 3aIII. vs. fIII. and c vs. g). The binding rate of microtubules and vimentin IFs increases to *r*_*b*,eff,Mg_ = 1.3·10^−2^ s^−1^. Generally, charged, suspended biopolymers in the presence of oppositely charged multivalent ions have been shown to attract these ions, leading to counterion condensation along the biopolymers. Consequently, the filaments attract each other through overscreening.^36,37^ Our data are in agreement with this effect. At high magnesium concentration, bonds form more likely and become stronger. Note that for both added magnesium chloride and TX100, the intermediate interactions (II) are decreased compared to the control conditions, due to a higher percentage of strong interactions (III) or weak interactions (I), respectively. Therefore, we conclude that both hydrophobic and electrostatic effects contribute to the direct interactions between microtubules and vimentin IFs.

When we moved the IF across the microtubule at an angle of 45° or horizontally in the direction of the microtubule (see Fig. 2e,f and Fig. 3h-k), we observed an increased binding rate (*r*_*b*,eff, 45°_ = 1.6 · 10^−2^ s^−1^ and *r*_*b*,eff,*x*_ = 2.4 · 10^−2^ s^−1^, respectively). This increase can be explained by an increased encounter rate of potential binding sites due to the different geometries (see Supplementary Information). Taking into account this geometric factor, we calculated the probability of a microtubule binding to a vimentin IF *p*_IF-MT_ for the different geometries and obtained *p*_IF-MT_ 6.1 · 10^−4^ per pair of vimentin unit-length filament and tubulin dimer, independent of the geometry. The breaking forces were found to be similar for the three different geometries (Fig. 3c, i and k). To test if vimentin IFs and microtubules co-align due to their interaction, as reported for migrating cells, ^18^ we relaxed the vimentin IF in the optical trap to allow for “zipping” events, or mixed the filaments in solution, but did not observe spontaneous bundling.

For a more profound understanding of the physical bond parameters, which are not accessible experimentally, we applied a modeling approach. We modeled the IF-microtubule interaction as a single molecular bond with force-dependent stochastic transitions between the bound and unbound state. The time-dependent force increase *F* (*t*) has an entropic stretching contribution^38,39^ for forces below 5 pN and increases linearly for higher forces as observed in the experiment.^40,41^ We assume that the binding (*b*) and unbinding (*u*) rates *r*_*b*_ and *r*_*u*_, respectively, depend on the applied force, an activation energy *E*_*Ab*_ or *E*_*Au*_, the thermal energy *k*_*B*_*T* and a distance *x*_*b*_ or *x*_*u*_ to the transition state, which is on the order of the distance between the IF and the microtubule at the site of the bond:

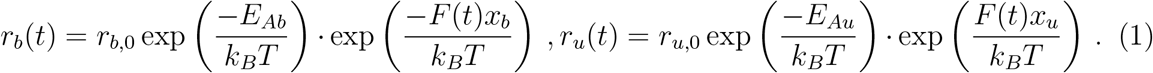

We summarize the force independent factor in Eq. (1) as an effective zero-force rate:

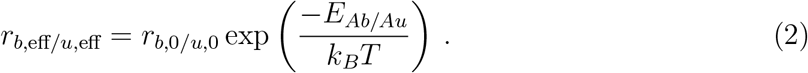

In contrast to the force and the effective binding rate *r*_*b*,eff_, neither *r*_*u*,eff_ nor *x*_*b*_ or *x*_*u*_ can be determined from our experimental data. Due to detailed balance, the sum *x*_*b*_ + *x*_*u*_ is constant.^42^ Since we only observed a small number of rebinding events under force, we focused on the unbinding processes and studied *x*_*u*_. Hence, we simulated IF-microtubule interactions for different sets of *r*_*u*,eff_ and *x*_*u*_ and compared the resulting distributions of breaking forces to our experimental data. We accepted the tested parameter sets if the distributions passed the Kolmogorov-Smirnov test with a significance level of 5%. The minimum and maximum of all accepted simulation results, shown as the borders of the green areas in Fig. 3c, e, g, I and k agree well with the experiments. Fig. 4a shows all accepted parameter pairs *r*_*u*,eff_ and *x*_*u*_ for the different buffer conditions (color code: gray (pure CB), orange (CB with TX100), blue (CB with additional magnesium); corresponding mixed colors for regions, where valid parameters overlap). Both parameters increase from additional magnesium (blue) across no addition (gray) to added TX100 (orange). A corresponding diagram for comparison of the different measuring geometries is shown in Supplementary Fig. 4. Whereas the force-free factor of the unbinding rate does not depend on the geometric configuration, the force-dependent factor is slightly more sensitive to force for a horizontal movement of the IF or a vertical movement of the IF with the microtubule turned by 45° than for a vertical movement of the IF perpendicular to the microtubule. To understand these data more intuitively, we calculated the energy diagrams, as plotted in Fig. 4b, using Eq. (8) (see Supplementary Information) considering the same buffer condition in unbound and bound (1 and 2) state or different buffer conditions (1 and 2) and the same state.

**Figure 4:**
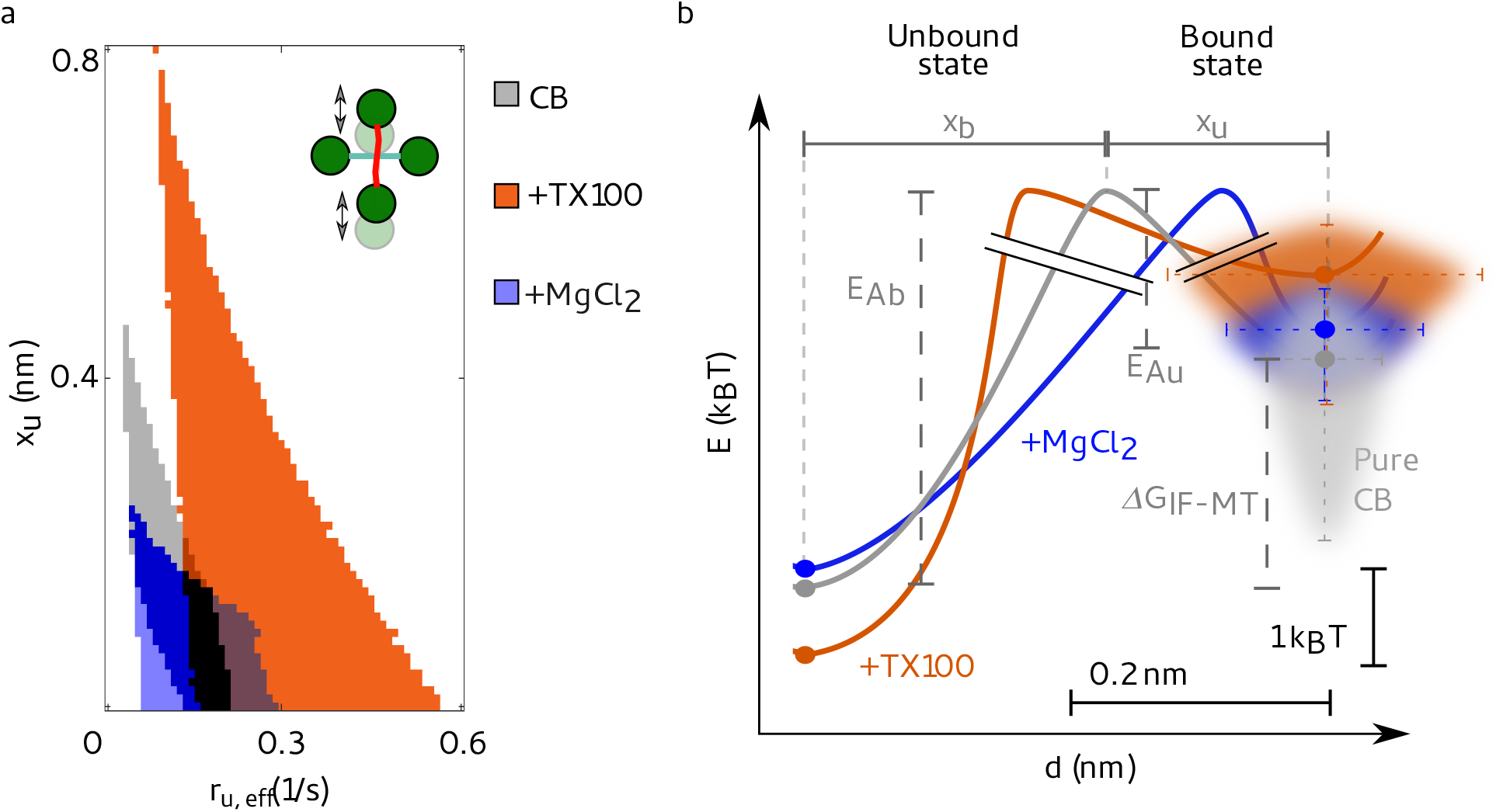
Hydrophobic and ionic reagents change the IF-microtubule bond properties. (a) Valid unbinding rates *r*_*u*,eff_ and potential widths *x*_*u*_ to simulate the experimental data shown in Fig. 3c, e and g for the different buffer conditions: pure CB (gray), CB with TX100 (orange) and CB with additional magnesium (blue). *r*_*u*,eff_ and *x*_*u*_ pairs which are valid for several buffer conditions are color coded by mixed colors. *r*_*u*,eff_ and *x*_*u*_ increase from additional magnesium chloride across pure CB to added TX100. (b) Energy landscape for the theoretical modeling of the IF-microtubule bond: A two-state model (unbound, bound) is sufficient to describe the experimental data shown in Fig. 3. From the binding and unbinding rates, we calculated the differences in activation energies, *E*_*Ab*_ and *E*_*Au*_, see Eq. (8) in the Supplementary Information, of bonds in different buffer conditions to open or close. However, the absolute values cannot be determined, as indicated by the graph break (black double-lines). For different buffer conditions, the position of the energy barrier relative to the unbound and bound state, *x*_*b*_ and *x*_*u*_, respectively, changes.

Surprisingly, both TX100 and additional magnesium only mildly affect the activation energies. Yet, for TX100 we observed a marked increase in distance to the transition state, *x*_*u*_ (compare Fig. 4b orange to gray), which we interpret as a “looser binding” between the IF and the microtubule. Thus the force-dependent term in Eq. (1) becomes more pronounced. TX100 can interact with hydrophobic residues and causes the filaments to stay further apart. Thus, the bond breaks at lower forces. Consequently, this further confirms that there is a hydrophobic contribution to the interactions in CB.

In contrast to TX100, magnesium strengthens the bond and keeps it closed even at higher forces as it is a divalent counterion between two negative charges. It lowers the distance to the transition state (compare Fig. 4b blue to gray) and the influence of the force-dependent term in Eq. (1). Hence, the opening of the bond depends less on the applied force compared to bonds in pure CB. Since CB already includes 2 mM magnesium, we assume that there is an electrostatic contribution to the interactions observed in CB as well.

We have shown that there are hydrophobic and electrostatic contributions to the interactions between IFs and microtubules and we have derived key parameters of these interactions by combining experiments with theoretical modeling. While we cannot exclude a steric contribution to the interaction, our measurements show that they are influenced by electrostatic and hydrophobic effects. We thus conclude that the interactions are modulated by electrostatics and hydrophobicity and directionally independent (see Supplementary Fig. 5 and Supplementary Movie 6). Furthermore, IFs assembled via dialysis are rather smooth,^43^ so that it is unlikely that their roughness causes interactions.

**Figure 5:**
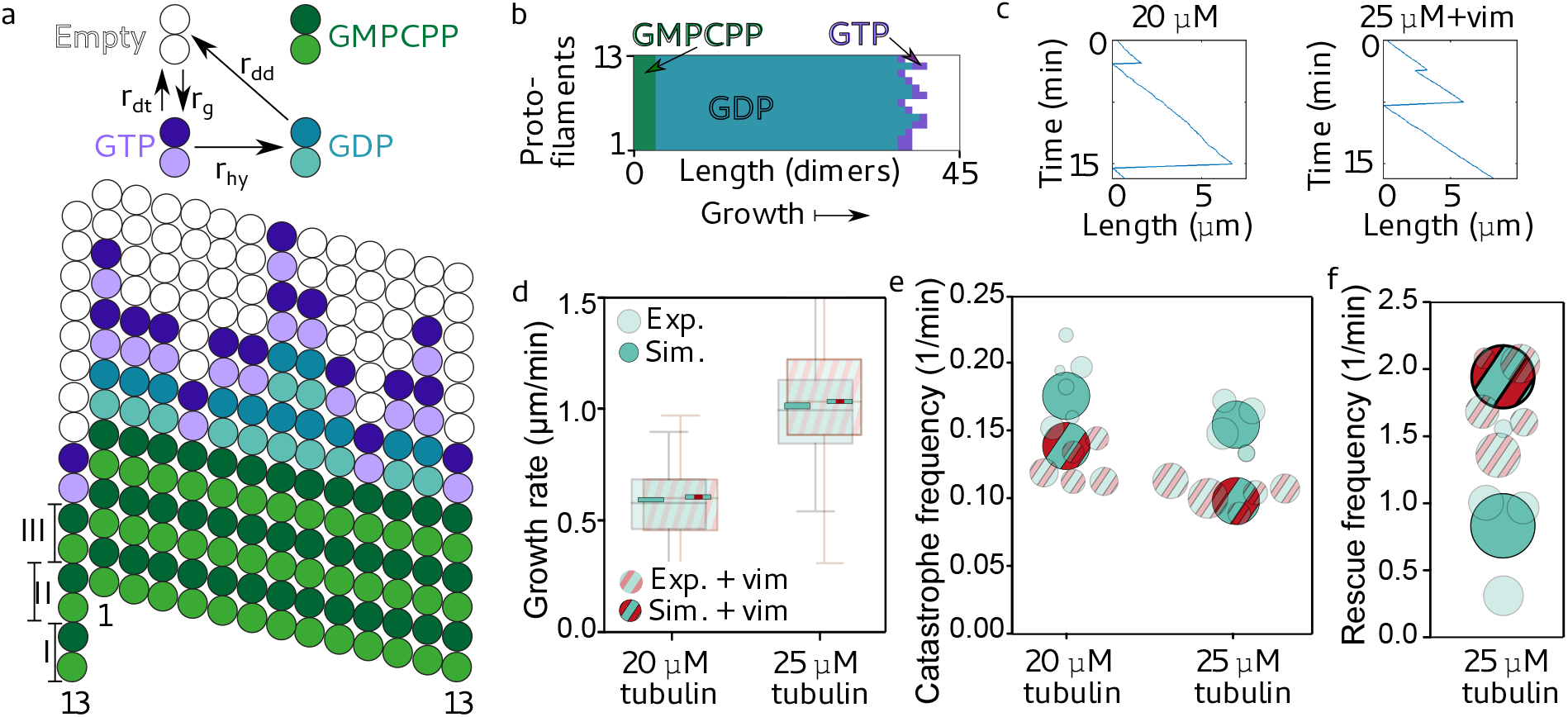
A Monte-Carlo simulation shows that transiently binding IFs stabilize dynamic microtubules. (a) Illustration of the reaction rates (top) and simulated microtubule lattice with 13 protofilaments and a seam with a longitudinal displacement of 1.5 dimers (bottom). (b) Typical simulated microtubule growing from a GMPCPP seed with dimers either in the GTP (purple) or GDP state (cyan). (c) Typical length-time plot (kymograph) of a simulated microtubule in 20 µM free tubulin solution without vimentin tetramers (left) or in 25 µM free tubulin solution with 2.3 µM vimentin tetramers (right). (d-f) We reproduced the experimental data shown in Fig. 1d-g (shown here in a semi-transparent fashion, for a vimentin concentration of 2.3 *µ*M) with our Monte-Carlo simulation (opaque). (d) Addition of vimentin neither changes experimental nor simulated microtubule growth rates at 20 or 25 µM. For clarity, the complete standard deviation of the experimental data is not shown, but presented in Fig. 1. (e) Addition of vimentin lowers the catastrophe frequency of dynamic microtubules for both tubulin concentrations studied here. (f) In case of 25 µM free tubulin, the rescue rate increases due to the stabilizing effect of surrounding vimentin IFs. The circle areas scale with the total microtubule depolymerization time as in the representation of the experimental data. All tubulin and vimentin concentrations are input concentrations.

To better understand how these interactions lead to the observed changes in microtubule dynamics, we again applied a modeling approach. We considered a microtubule as a dynamic lattice with GTP and GPD dimers^20,26^ as sketched in Fig. 5a. The lattice consists of 13 protofilaments and has a seam between the first and 13th protofilament. We describe the microtubule dynamics by three reactions: (i) A GTP dimer associates with a rate *r*_*g*_, (ii) a GTP dimer is hydrolyzed with a rate *r*_*hy*_, or (iii) a GDP or GTP dimer dissociates with a rate *r*_*dd*_ or *r*_*dt*_, respectively, depending on the number of neighboring dimers, see Eq.(12) in the Supplementary Information. A snapshot of the simulated microtubule during growth is shown in Fig. 5b. With a Monte-Carlo simulation, we obtained typical simulated kymographs (Fig. 5c). As for the experiments (semi-transparent data in Fig. 5d-f), we determined the growth rate, the catastrophe and the rescue frequency from the simulations (opaque in Fig. 5d-f).

To simulate microtubules in the presence of vimentin IFs, we considered the interactions of IFs and microtubules measured with OT experiments again. Assuming that IFs bind stochastically to the microtubule lattice, we estimated binding and unbinding rates from the OT experiments (see the Supplementary Information for a detailed description of the model). Based on these rates, we calculated the microtubule-IF binding energy (see Eq. (9) in the Supplementary Information) to be ∆*G*_IF-MT_ = 2.3 *k*_*B*_*T* as sketched in Fig. 4b. Thus, IF binding stabilizes the binding of tubulin dimers in the microtubule lattice by 2.3 *k*_*B*_*T*. The additional binding energy can be interpreted as a direct increase of the total binding energy of the respective tubulin dimer or as an increased longitudinal binding energy to the next tubulin dimer. Our experiments do not resolve the precise molecular interaction mechanism, such as cross-linking of neighboring tubulin dimers or structural changes in the tubulin dimers upon binding of a vimentin IF. Likewise, we cannot distinguish whether the vimentin IF is bound to a single tubulin dimer or to multiple dimers. However, our coarse description approach includes all these different scenarios. Specifically, in case of a bond involving multiple dimers, unbinding from these dimers must be cooperative since we do not observe step-wise unbinding in optical trapping experiments. Such cooperativity does not change the total energy required for unbinding. In agreement with our experimental data, the transient binding of IFs leaves the growth rate unaffected. Intriguingly, we observed that IF binding to tubulin dimers in the lattice reduces the catastrophe frequency. The increased binding energy of a dimer also raises the rescue frequency. These results are in striking agreement with our observation in TIRF experiments, while the only additional input to the simulation that includes the surrounding vimentin IFs are the parameters from OT experiments. Thus, stochastic, transient binding of IFs to microtubules as in the OT experiments is sufficient to explain the observed changes in microtubule dynamics in presence of IFs.

By combining the results from OT and TIRF experiments, we estimated the total binding energy of a tubulin dimer within the lattice at the microtubule tip before catastrophe. From the IF-microtubule bond breaking events in the OT experiments, including the corresponding simulations, we calculated the IF-microtubule bond energy ∆*G*_IF-MT_ = 2.3 *k*_*B*_*T* and the unbinding rate *r*_*u*,eff_ of microtubules and vimentin IFs (Fig. 6a). From the TIRF experiments, we determined the catastrophe frequency *f*_cat,IF-MT_ of a microtubule bound to a vimentin IF. At the beginning of the catastrophe, a vimentin IF unbinds from the tubulin dimer, so that the energy ∆*G*_IF-MT_ is released. Simultaneously, the dimer depolymerizes from the lattice and the energy ∆*G*_*tb*_ is set free (Fig. 6b). The only additional energy released during microtubule catastrophe in TIRF experiments compared to the OT experiments is the binding energy to the surrounding tubulin dimers (Fig. 6c). Thus, comparing the rates of IF-microtubule unbinding and microtubule catastrophe during binding to a vimentin IF as in Eq. (15) in the Supplementary Information, results in an estimation of the average tubulin binding energy ∆*G*_*tb*_ between 5.7 *k*_*B*_*T* and 7.2 *k*_*B*_*T* in the lattice at the tip. These values for ∆*G*_*tb*_ are on the order of magnitude expected from interferometric scattering microscopy and from computational studies, although slightly lower, possibly due to different buffer conditions.^20,27,44^ Our combination of experiments provides a new way of determining such binding energies and may, from a broader perspective, be generally applied to proteins which bind to microtubules.

**Figure 6:**
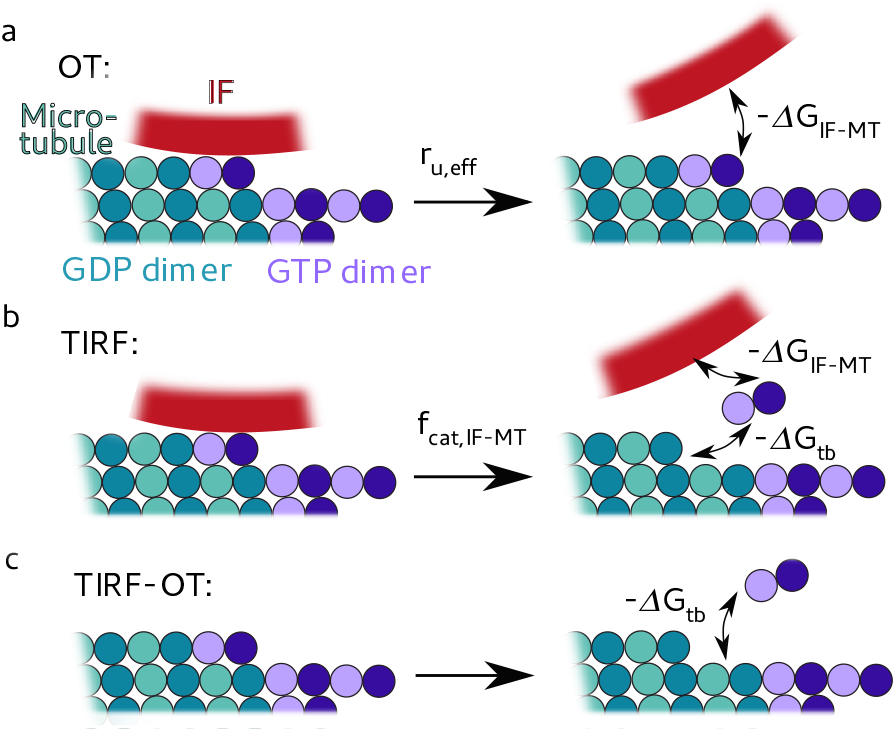
We estimated the binding energy of a single tubulin dimer by combining our OT and TIRF experiments. (a) From the OT experiments including the simulations, we determined the unbinding rate of microtubules and vimentin IFs *r*_*u*,eff_ and the released energy −∆*G*_IF-MT_ during unbinding. (b) In TIRF experiments, a tubulin dimer dissociates from the microtubule and the vimentin IF, so that the total energy −∆*G*_*tb*_ −∆*G*_IF-MT_ is released. We calculated the catastrophe frequency of a microtubule *f*_cat,IF-MT_ in case a vimentin IF is bound to the microtubule. (c) We estimated the average binding energy ∆*G*_*tb*_ of a tubulin dimer in the microtubule lattice before catastrophe by subtracting the released energies from (b) (−∆*G*_*tb*_ −∆*G*_IF-MT_) and by dividing the unbinding rate *r*_*u*,eff_ of microtubules and vimentin IFs by the catastrophe frequency *f*_cat,IF-MT_ of microtubules bound to a vimentin IF as in Eq. (15) in the Supplementary Information.

Our study examined interactions between microtubules and vimentin IFs. We showed that vimentin IFs stabilize microtubules by direct interactions, which is in strong contrast to previous findings, ^14^ where, however, only interactions between microtubules and short IF peptides were considered. Whereas the microtubule growth rate remains unchanged, the stabilization by vimentin IFs leads to a reduction in the catastrophe frequency and increased rescue of depolymerizing microtubules. We pinpoint the source of this stabilizing effect to a stochastic, transient binding of IFs to microtubules by directly measuring the interactions of single filaments. Both hydrophobic and electrostatic effects are involved in bond formation. The presence of cations likely contributes to the attractive interactions between the negatively charged filaments. The buffer in which we conducted the measurements contained potassium and magnesium, two of the most abundant cations in cells.^45^ The free magnesium concentration is on the order of a few mM in most mammalian cells, ^46^ similar to our experiments. Magnesium ions have been previously described to cross-link vimentin IFs,^47–51^ and we showed that they can modulate the IF-microtubule bond strength. Since our magnesium concentrations were close to physiological values, the magnesium induced IF-microtubule binding we observed may occur in cells as well. Therefore, although molecular motors and cross-linkers contribute to establishing links between IFs and microtubules in cells, our results indicate that more fundamental, direct attractive interactions may also participate in the crosstalk of the two cytoskeletal subsystems in cells. The interactions we observed might thus be linked to cytoskeletal crosstalk between microtubules and vimentin IFs in living cells^18^ and the coexistensive network formation in migrating astrocytes.^16^

Gan et al.^18^ reported the stabilization of the microtubule network by vimentin IFs. Complementary to this finding, we showed microtubule stabilization by transient interactions with IFs on the single filament level without coalignment or bundling of the two filament types. We found that vimentin IFs and microtubules do not spontaneously coalign and that bundling is unlikely, which indicates that the cell has to activate additional interaction mechanisms, e.g. via proteins, to induce the coalignment in migrating cells. Such mechanisms would allow the cell to switch to a migrating state in a controlled way. Moreover, our results indicate that by increasing the local vimentin filament concentration by transport of filament fragments along microtubules,^52^the cell can locally tune the dynamic instability of microtubules, which might be a vital mechanism for cell polarization and preparation for migration. The rapid, but unfrequent binding of IFs and microtubules we observed suggests that only certain microtubule and IF subunits can bind. Thus, we hypothesize that controlling which subunits can bind (e.g. by posttranslational modifications) may provide another path for the cell to regulate the stabilization of microtubules by IFs.

There is growing evidence that a mechanical coupling between the cytoskeletal subsystems is necessary for many cellular functions such as polarization, migration and mechanical resistance.^2,53,54^ In particular, vimentin deficient cells exhibit a less robust microtubule network orientation^18^ and stronger microtubule fluctuations,^17^ and they show impaired migration, contractility and resistance to mechanical stress. ^55–57^ Therefore, future research might help to explore the implications of our findings for cell mechanics and function. Furthermore, our study fosters understanding of emergent material properties of hybrid networks composed of cytoskeletal filaments and provides a basis for interpreting rheology data. Our combination of experiments also offers a new approach to estimate the tubulin bond energy within the microtubule lattice, which is a vital parameter to understand microtubule dynamics, mechanics and function.^20–27^

## Materials and Methods

### Vimentin purification, labeling and assembly

Vimentin C328N with two additional glycines and one additional cysteine at the C-terminus was recombinantly expressed as previously described^58–60^ and stored at −80°C in 1 mM EDTA, 0.1 mM EGTA, 0.01 M MAC, 8 M urea, 0.15-0.25 M potassium chloride and 5 mM TRIS at pH 7.5.^60^ After thawing, we labeled the vimentin monomers with the fluorescent dye ATTO647N (AD 647N-41, AttoTech, Siegen, Germany) and with biotin via malemeide (B1267-25MG, Jena Bioscience, Jena, Germany) as described in Refs. 61–63. We mixed labeled and unlabeled vimentin monomers, so that in total 4% of all monomers were fluorescently labeled, a maximum of 20 % was biotin labeled and all other monomers were unlabeled.^59,63^ We reconstituted vimentin tetramers by first dialyzing the protein against 6 M urea in 50 mM phoshate buffer (PB), pH 7.5, and then in a stepwise manner against 0 M urea (4, 2, 0 M urea) in 2 mM PB, pH 7.5,^64^ followed by an additional dialysis step against 0 M urea, 2 mM PB, pH 7.5, overnight at 10°C. To assemble vimentin into filaments, we dialyzed the protein into assembly buffer, i.e. 100 mM KCl, 2 mM PB, pH 7.5, at 36°C overnight.^60,64^

### Tubulin purification and labeling

We purified tubulin from fresh bovine brain by a total of three cycles of temperature-dependent assembly and disassembly in Brinkley buffer 80 (BRB80 buffer; 80 mM PIPES, 1 mM EGTA, 1 mM MgCl_2_, pH 6.8, plus 1 mM GTP) as described in Ref. 65. After two cycles of polymerization and depolymerization, we obtained microtubule-associated protein (MAP)-free neurotubulin by cation-exchange chromatography (1.16882, EMD SO_3_, Merck, Darmstadt, Germany) in 50 mM PIPES, pH 6.8, supplemented with 0.2 mM MgCl_2_ and 1 mM EGTA.^66^We prepared fluorescent tubulin (ATTO488- and ATTO565-labeled tubulin; AttoTech AD488-35 and AD565-35, AttoTech) and biotinylated tubulin (NHS-biotin, 21338, Thermo Scientific, Waltham, MA, USA) as previously described.^67^ In brief, microtubules were polymerized from neurotubulin at 37°C for 30 min, layered onto cushions of 0.1 M NaHEPES, pH 8.6, 1 mM MgCl_2_, 1 mM EGTA 60% v/v glycerol, and sedimented by centrifugation at 250,000 × g 37°C for 1 h. We resuspended microtubules in 0.1 M Na-HEPES, pH 8.6, 1 mM MgCl_2_,1 mM EGTA, 40 % v/v glycerol and labeled the protein by adding 1/10 volume 100 mM NHS-ATTO or NHS-biotin for 10 min at 37°C. We stopped the labeling reaction by adding 2 volumes of BRB80×2, containing 100 mM potassium glutamate and 40% v/v glycerol. Afterwards, we centrifuged microtubules through cushions of BRB80 containing 60% v/v glycerol. We resuspended microtubules in cold BRB80 and performed an additional cycle of polymerization and depolymerization before we snap-froze the tubulin in liquid nitrogen and stored it in liquid nitrogen until use.

### Microtubule seeds for TIRF experiments

We prepared microtubule seeds at 10 µM tubulin concentration (30% ATTO-565-labeled tubulin and 70% biotinylated tubulin) in BRB80 supplemented with 0.5 mM GMPCPP at 37°C for 1 h. We incubated the seeds with 1 µM taxol for 30 min at room temperature and then sedimented them by centrifugation at 100,000 × g for 10 min at 37°C. We discarded the supernatant and carefully resuspended the pellet in warm BRB80 supplemented with 0.5 mM GMPCPP and 1 µM taxol. We either used seeds directly or snap-froze them in liquid nitrogen and stored them in liquid nitrogen until use.

### Sample preparation for optical trapping experiments

We prepared stabilized microtubules with biotinylated ends for OT by first polymerizing the central part of the microtubules through step-wise increase of the tubulin concentration. Initially, a 3 µM tubulin (5% ATTO-488-labeled) solution in M2B buffer (BRB80 buffer supplemented with 1 mM MgCl_2_) in the presence of 1 mM GMPCPP (NU-405L, Jena Bio-science) was prepared at 37°C to nucleate short microtubule seeds. Next, the concentration was increased to a total of 9 µM tubulin in order to grow long microtubules. To avoid further microtubule nucleation, we added 1 µM tubulin at a time from a 42 µM stock solution (5% ATTO-488-labeled) and waited 15 min between the successive steps. To grow biotinylated ends, we added a mix of 90% biotinylated and 10% ATTO-565-labeled tubulin in steps of 0.5 µM from a 42 µM stock solution up to a total tubulin concentration of 15 µM. We centrifuged polymerized microtubules for 10 min at 13000 × g to remove any non-polymerized tubulin and short microtubules. We discarded the supernatant and carefully resuspended the pellet in 800 µL M2B-taxol (M2B buffer supplemented with 10 µM taxol (T7402, Merck)). By keeping the central part of the microtubules biotin-free (see color code in Fig. 2a: biotin-free microtubule in cyan and biotin labeled microtubule ends in green), we ensured that any streptavidin molecules detaching from the beads could not affect interaction measurements by cross-linking the filaments.

For measurements in the microfluidic chip by OT, we prepared four solutions for the four different microfluidic channels as sketched in Fig. 2a: (I) We diluted streptavidin-coated beads with an average diameter of 4.5 µm (PC-S-4.0, Kisker, Steinfurt, Germany) 1:83 with vimentin assembly buffer. (II) We diluted the vimentin IFs 1:667 with vimentin assembly buffer. (III.) We diluted the resuspended microtubules 1:333 with CB. (IV.) We combined suitable buffer conditions for microtubules and for vimentin IFs, respectively, to a combination buffer (CB) containing 1 mM EGTA (03777, Merck), 2 mM magnesium chloride, 25 mM PIPES (9156.4, Carl Roth, Karlsruhe, Germany), 60 mM potassium chloride (6781.3, Carl Roth) and 2 mM sodium phosphate (T879.1 and 4984.2, Carl Roth) at pH 7.5. We included an oxygen scavenging system consisting of 1.2 mg/mL glucose (G7528, Merck), 0.04 mg/mL glucose oxidase (G6125-10KU, Merck), 0.008 mg/mL catalase (C9322-1G, Merck) and 20 mM DTT (6908.2, Carl Roth). Additional 0.01 mM taxol (T1912-1MG, Merck) stabilized the microtubules. For measurements with TX100, we added 0.1 % (w/v) Triton-X 100 (TX100; 3051.3, Carl Roth) and in case of measurements with a total magnesium concentration of 20 mM, we added 18 mM MgCl_2_. We filtered the solutions through a cellulose acetate membrane filter with a pore size of 0.2 µm (7699822, Th. Geyer, Renningen, Germany).

### Optical trapping experiment

We performed the OT experiments using a commercial setup (C-Trap, LUMICKS, Amsterdam, The Netherlands) equipped with quadruple optical tweezers, a microfluidic chip and a confocal microscope. Beads, microtubules, measuring buffer and IFs were flushed into four inlets of the microfluidic chip as sketched in Fig. 2a. For each measurement, four beads were captured and then calibrated in the buffer channel using the thermal noise spectrum. One bead pair (beads 1 and 2) was moved to the vimentin IF channel and incubated there until a filament bound to beads (Fig. 2aII.). Meanwhile, the other bead pair (beads 3 and 4) was kept in the measuring buffer channel, so that no filaments adhered to those beads. To capture a microtubule (Fig. 2aIII.), beads 3 and 4 were moved to the microtubule channel, while bead 1 and 2 stayed in the measuring buffer channel. Once a microtubule was bound to beads 3 and 4 and an IF to beads 1 and 2, the bead pair with the IF was horizontally turned by 90°(Supplementary Fig. 3a) and moved up in *z*-direction by 4.9 µm. The bead pair holding the IF was moved in the *x*-*y*-plane so that the central part of the IF was positioned above the center of the microtubule (Fig. 2aIV. and Supplementary Fig. 3a). To bring the IF and microtubule into contact, the IF was moved down in *z*-direction until the microtubule was pushed into focus or slightly out of focus. In a portion of the experiments, we turned the microtubule by 45°and visually controlled the angle by fluorescence microscopy.

The IF was moved perpendicularly to the microtubule in the *y*-direction in the *x*-*y*-plane at 0.55 µm/s, while we measured the forces in the *x*- and *y*-direction on bead 1. For a horizontal movement, we moved the IF perpendicularly to the microtubule in the *x*-direction in the *x*-*y*-plane at 0.55 µm/s. Simultaneously, we recorded confocal images to see whether an interaction occured. In case no interaction occured after two movements in the *x*-*y*-plane, the IF was moved down in *z*-direction by 0.4 µm and the movement in the *x*-*y*-plane was repeated. The experiment ended when the microtubule broke off the bead, or the IF or microtubule broke. In case of the vertical movement in a perpendicular configuration, we measured 57 pairs of microtubules and vimentin IFs in CB, 38 pairs with TX100 and 36 pairs with additional magnesium chloride. In total, we moved the IFs 744 times vertically and perpendicularly to the microtubules in CB, 704 times in CB with TX100 and 542 times in CB with additional magnesium chloride. In the case of the 45°-configuration we studied 49 pairs of microtubules and vimentin IFs and completed 467 movements. In the case of horizontal movement of the IF in perpendicular configuration with respect to the microtubule, we studied 43 filament pairs and completed 504 movements.

### Optical trapping data analysis

The OT data were processed with self-written Matlab (MathWorks, Natick, MA, USA) scripts. In case of the vimentin IF movement in the *y*-direction and a perpendicular orientation to the microtubule, we analyzed the component of the force *F*_1*y*_ acting on bead 1 in the *y*-direction for each filament pair, since the forces in *x*-direction were balanced by the IF, as sketched in Supplementary Fig. 3b. From the raw force data, we manually selected the force data containing an interaction. Due to interactions of the energy potentials of the different traps, some data sets exhibited a linear offset which we subtracted from the data. From the interaction-free force data, we determined the experimental error by calculating the standard deviation in the force of the first 20 data points. We defined an interaction as soon as the force *F*_1*y*_ as shown in Fig. 2d deviated by more than 5*σ*_*F*_ from the mean of the first 20 data points, where *σ*_*F*_ is the standard deviation of the force without interactions in each data set. Typically, the force increased as shown in Fig. 2d until the interaction ended with a fast force decrease as marked by ∆*F*_1*y*_. We did not take breaking forces below 0.5 pN into account because they may be caused by force fluctuations. Since the force detection of trap 1 is the most accurate one in the setup, we analyzed the force on bead 1 only. To determine the total breaking force *F*_*B*_, we multiplied the force *F*_1*y*_ acting on bead 1 in *y*-direction with a correction factor *c*_*F*_ that is based on the geometry of the experiment. *c*_*F*_ depends on the distance between bead 1 and 2 *d*_MT_ and the distance *d*_IF-MT_ from bead 1 to the contact point of the IF and the microtubule as sketched in Supplementary Fig. 3c:

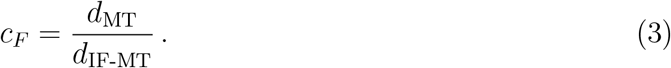

For the total force *F*_*C*_ acting on the IF-microtubule bond, we get:

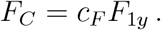

Thus, when an IF-microtubule bond breaks, the total force difference *F*_*B*_ is:

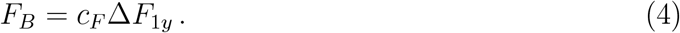

In case of a 45°angle between microtubule and vimentin IF and in case of IF movement in the *x*-direction the force data were analyzed in the same way as described above with the following differences:

For the case, where the microtubule was turned by 45°, we calculated the geometric factor in two different ways, depending on the geometry at the moment of bond breakage: (i) If the bond breaks at a higher *y*-position than bead 2 (see Supplementary Fig. 3d), we assume that the total force acting on the IF-microtubule bond is measured by the complete force *F*_1_ acting on bead 1. Thus, *c*_*F*_ = 1, and the breaking force is:

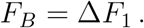

(ii) If the bond breaks at a lower *y*-position than bead 2 (see Supplementary Fig. 3e), the geometric factor is:

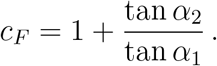

We analyzed *F*_1*y*_ again and the breaking force was calculated with Eq. (4).

In case the IF was moved perpendicularly to the microtubule in the *x*-direction, we analyzed the force *F*_1*x*_ acting on bead 1 in the *x*-direction. We did not need to correct for the geometry of the experiment, thus, *c*_*F*_ = 1:

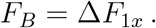

### Preparation of passivated cover glasses for TIRF experiments

We cleaned cover glasses (26 × 76 mm^2^, no. 1, Thermo Scientific) by successive chemical treatments: (i) We incubated the cover glasses for 30 min in acetone and then (ii) for 15 min in ethanol (96% denatured, 84836.360, VWR, Radnor, PA, USA), (iii) rinsed them with ultrapure water, (iv) left them for 2 h in Hellmanex III (2% (v/v) in water (Hellma Analytics, Müllheim, Germany), and (v) rinsed them with ultrapure water. Subsequently, we dried the cover glasses using nitrogen gas flow and incubated them for three days in a 1 g/L solution of 1:10 silane-PEG-biotin (PJK-1919, Creative PEG Works, Chapel Hill, NC, USA) and silane-PEG (30 kDa, PSB-2014, Creative PEG Works) in 96% ethanol and 0.02% v/v hydrochloric acid, with gentle agitation at room temperature. We subsequently washed the cover glasses in ethanol and ultrapure water, dried them with nitrogen gas and stored them at 4°C for a maximum of four weeks.

### TIRF experiments

We used an inverted microscope (IX71, Olympus, Hamburg, Germany) in TIRF mode equipped with a 488-nm laser (06-MLD, 240 mW, COBOLT, Solna, Sweden), a 561-nm laser (06-DPL, 100 mW, COBOLT) and an oil immersion TIRF objective (NA = 1.45, 150X, Olympus). We observed microtubule dynamics by taking an image every 5 s for 15 – 45 min using the CellSense software (Olympus) and a digital CMOS camera (ORCA-Flash4.0, Hamamatsu Photonics, Hamamatsu, Japan).

For TIRF experiments, we built flow chambers from passivated cover glasses and double sided tape (70 µm height, 0000P70PC3003, LiMA, Couzeix, France). We flushed 50 µg/mL neutravidin (A-2666, Invitrogen, Carlsbad, CA, USA) in BRB80 into the chamber and incubated for 30 s. To remove free neutravidin, we washed with BRB80. Afterwards, we flushed microtubule seeds diluted 300 × in BRB80 into the chamber and incubated for 1 min before we removed free floating seeds by washing with BRB80 supplemented with 1% BSA. Then, a mix containing 0.5 mg/mL or 0.8 mg/mL (corresponding to 2.34 µM or 3.74 µM) vimentin tetramers (4% ATTO-565-labeled), 20 µM or 25 µM tubulin dimers (20% ATTO488-labelled), 0.65% BSA, 0.09% methyl cellulose, 2 mM phosphate buffer, 2 mM MgCl_2_, 25 mM PIPES, 1 mM EGTA, 60 mM KCl, 20 mM DTT, 1.2 mg/mL glucose, 8 µg/mL catalase and 40 µg/mL glucose oxidase, pH 7.5, was perfused into the chamber. To avoid evaporation and convective flow, we closed the chamber with vacuum grease and placed it on the stage of the TIRF microscope that was kept at 37°C.

From the TIRF movies, kymographs were created using the reslice function of ImageJ (ImageJ V, version 2.0.0-rc-69/1.52p). From the kymographs, microtubule growth and depolymerization velocities, catastrophe and rescue frequencies were estimated. We calculated the catastrophe frequency for each experiment as

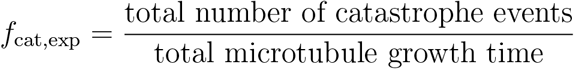

and the rescue frequency as

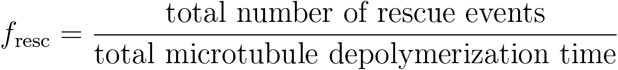

The total growth time was 800-2000 min per condition, the total depolymerization time 50-70 min per condition. The growth rate plot in Fig. 1d contains between 106 and 163 measurements per condition.

### Determination of vimentin filament length distributions

To measure the lengths of vimentin filaments (see Supplementary Fig. 1), we prepared five 1.5 mL reaction tubes with 15 µL of a mix of 2.3 µM vimentin in CB including all additions as used for the TIRF experiments such as methyl cellulose, GTP and oxygen scavenger (see previous section for the exact composition of the buffer). We then incubated the mix at 37°C for 5, 10, 20, 30 or 45 min. The filament assembly was stopped by adding 25 volumes of buffer to the tubes. 5 µL of each diluted mix were then pipetted on a cover glass and a second cover glass was placed on top. Images were taken with an inverted microscope (IX81, Olympus) using the CellSense software (Olympus), a 60x oil-immersion PlanApoN objective (Olympus) and an ORCA-Flash 4.0 camera (Hamamatsu Photonics). The filament lengths were determined using the semi-automated JFilament 2D plugin (Lehigh University, Bethlehem, PA, USA, version 1.02) for ImageJ (version 2.0.0-rc-69/1.52p).

## Modeling

A detailed description of the modeling approaches is provided in the Supplementary Information.

In brief, breaking of microtubule-IF interactions was modeled as a force-dependent two-state model and distributions of breaking forces were determined both by stochastic simulations and by numerically solving for the first passage time distribution. The time-dependent force accounts for the pulling speed and the entropic elasticity of the filament. The unknown parameters were varied systematically to find regions in parameter space consistent with the OT experiments for different buffer conditions and different measurement geometries.

The simulation of dynamic microtubules was adapted from Refs. 20,26 by additionally considering the binding energy of a microtubule to a vimentin IF obtained from the OT experiments. We thus simulated dynamic microtubules embedded in a vimentin IF network. The binding energy of a microtubule dimer in the lattice at the tip of a microtubule was calculated from an energy balance comparing the unbinding rate of microtubules and vimentin IFs, and the catastrophe frequency with and without vimentin.

## Acknowledgements

We thank Thomas Wenninger, Helge Schmidt and Sarah Adio for their support concerning the construction of the TIRF setup. We are grateful for purified tubulin and vimentin from Manuel Théry, Laurent Blanchoin, Jérémie Gaillard, Susanne Bauch. We thank Susanne Bauch, Susanne Hengst and Ulrike Schulz for technical support. The work was financially supported by the European Research Council (ERC, Grant No. CoG 724932, to S.Kö.), the European Molecular Biology Organization (Long Term Fellowship No. 1164-2018, to L.S.) and the Studienstiftung des deutschen Volkes e.V. (fellowship to C.L.) This research was conducted within the Max Planck School Matter to Life (to S.Kö. and S. Kl.) supported by the German Federal Ministry of Education and Research (BMBF) in collaboration with the Max Planck Society. The work further received financial support via an Excellence Fellowship of the International Max Planck Research School for Physics of Biological and Complex Systems (IMPRS PBCS, fellowship to A.V.S.).

## Author Contributions

S.Kö. conceived and supervised the project. S.Kö. and L.S. designed the experiments. L.S. and C.L. performed all experiments and analyzed the data. A.V.S. helped performing the quadruple optical trap experiments. C.L. and S.Kl. designed and performed numerical simulations. All authors contributed to writing the manuscript.

## Data availability

The data that support the findings of this study are available from the corresponding authors upon reasonable request.

## Code availability

The source codes of the numerical simulations are part of the Supplementary Information. The codes for the analysis of the model simulations are available from the corresponding authors upon request.

