## Supplementary Information for "Vimentin Intermediate Filaments Stabilize Dynamic Microtubules by Direct Interactions"

§*Max Planck School "Matter to Life"*

### Supplementary Figures

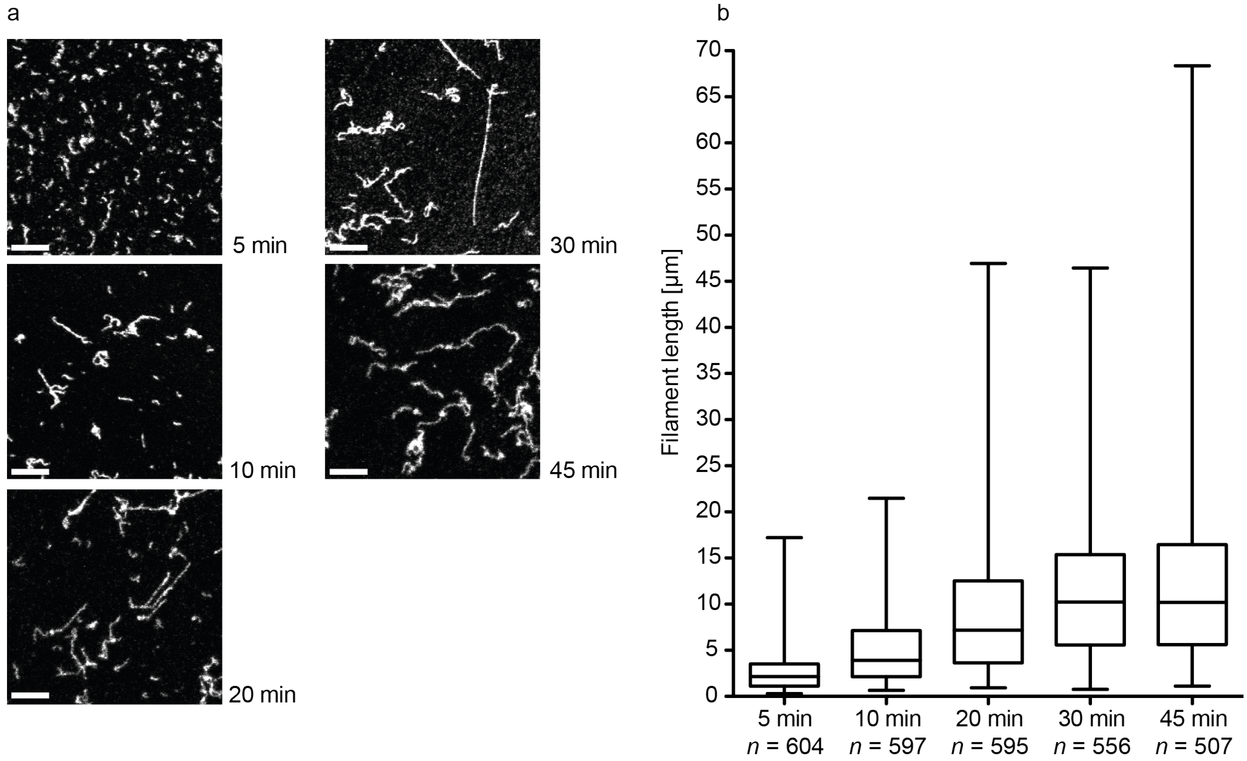


Figure 1: Temporal evolution of vimentin filament length in CB at a concentration of 2.3 µM. (a) Typical epifluorescence microscopy images at different time points after start of assembly by CB buffer addition. Scale bars correspond to 5 $\mu$m. We started the TIRF measurements 5-10 mins after initiation of the assembly and they ran for 30 to 45 mins. (b) Traced filament lengths at different time points after starting the assembly. $n$ is the number of filaments traced. Boxplots include the median as the center line, the 25th and 75th percentiles as box limits and the entire data range as whiskers.

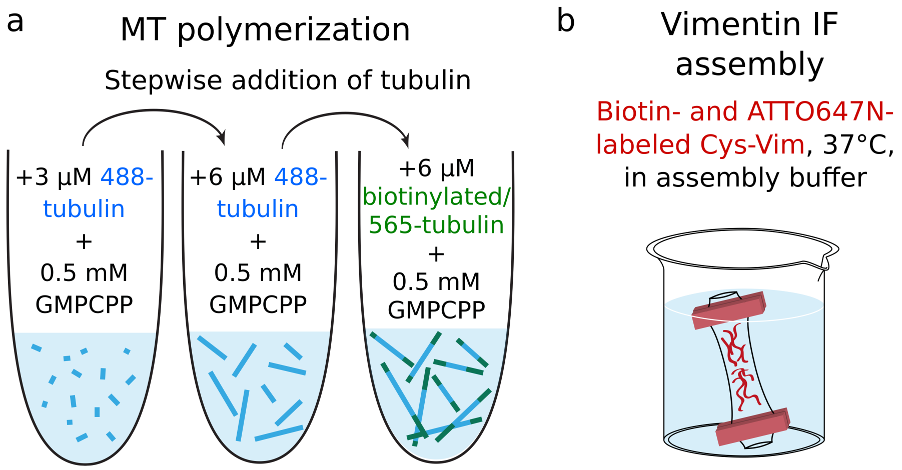


Figure 2: (a) Schematic of microtubule preparation. GMPCPP stabilized microtubules were prepared by first growing the central, biotin-free part through stepwise addition of ATTO488 labeled tubulin (blue). Biotinylated ends were added by stepwise addition of ATTO565 labeled and biotinylated tubulin (green). (b) Schematic of IF preparation. Biotinylated and ATTO647N labeled vimentin protein was assembled into filaments overnight via dialysis into assembly buffer.

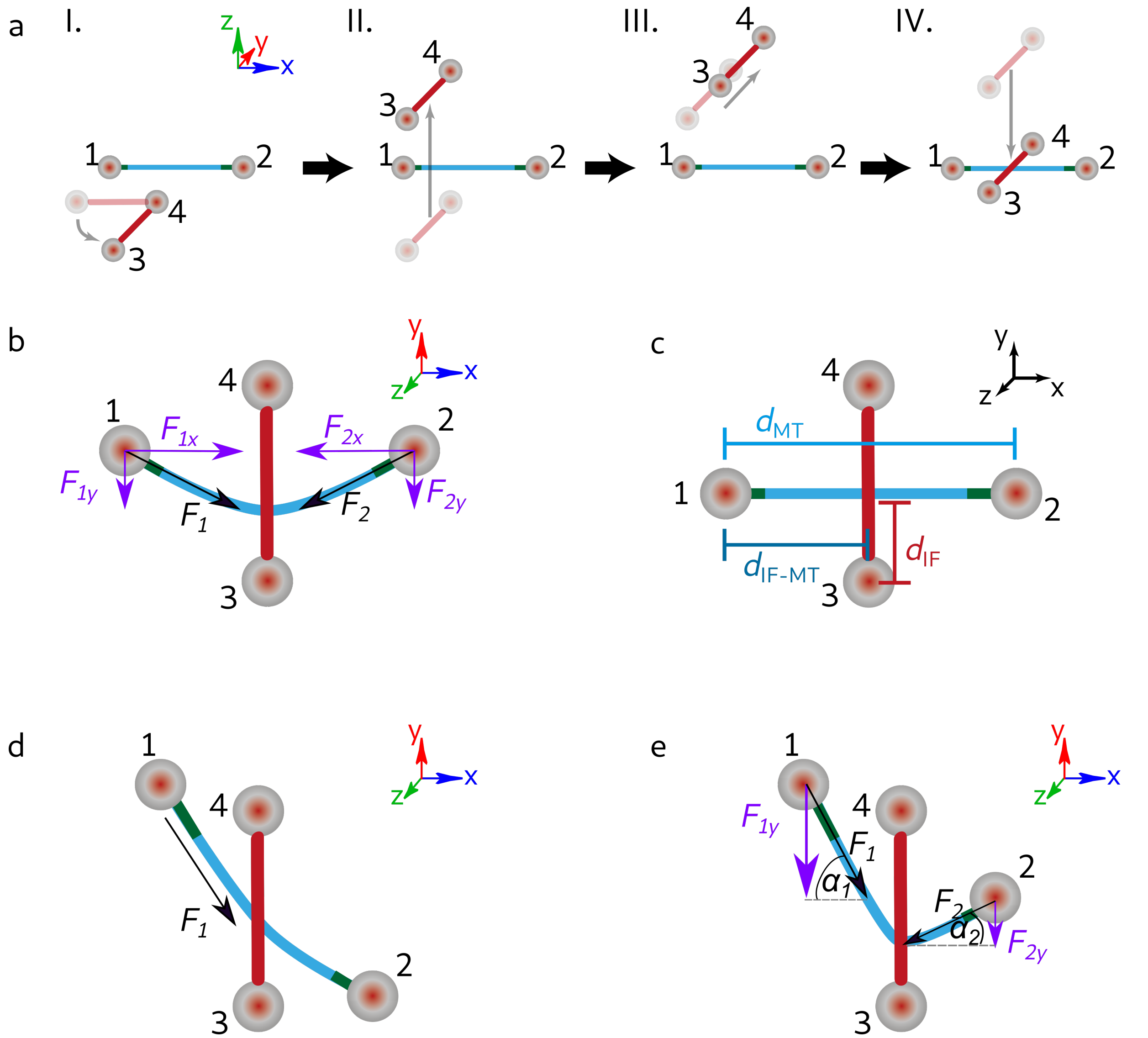


Figure 3: Protocol and geometry of the OT experiments. (a) To measure direct interactions between a single IF (red) and a single microtubule (blue), we turned the vimentin filament in the $x$-$y$ plane so that it was perpendicularly aligned to the microtubule (MT in $x$-direction, IF in $y$-direction (I). We moved the IF upwards in $z$-direction (II.) and moved it in the $x$-$y$ plane so that the center of the IF was positioned above the center of the microtubule (III.). Next (IV.), we moved the IF downwards in $z$-direction until it was in the same $x$-$y$ plane as the microtubule; the IF and microtubule were then in contact. (b) Forces acting on the IF and the microtubule. (c) Definition of the length scales required for the analysis of the OT data. (d) Geometric configuration and analyzed force if the vimentin IF was moved in the $y$-direction, the microtubule was turned by 45°, and the point of interaction was located higher in $y$ than bead 2. (e) Definition of forces and angles for the same configuration as in d if the point of interaction was located lower in $y$ than bead 2.

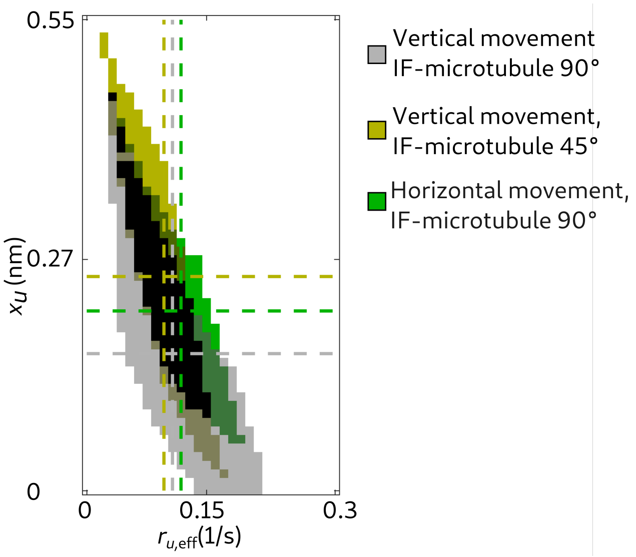


Figure 4: Valid unbinding rates $r_{u,\text{eff}}$ and potential widths $x_u$ to simulate the experimental data shown in Fig. 3c, i and k for the different geometric configurations of the OT experiment (all in pure CB): vertical movement of the IF perpendicular to the microtubule (gray), vertical movement of the IF with the microtubule turned by 45° (yellow), and horizontal movement of the IF perpendicular to the microtubule (green). $r_{u,\text{eff}}$ and $x_u$ pairs, which are valid for several geometric configurations, are color coded by mixed colors. The centroid positions of the areas are projected on the $r_{u,\text{eff}}$ and $x_u$ axes shown by dashed lines in the corresponding colors. These are the mean values for $r_{u,\text{eff}}$ and $x_u$ we used for further calculations. The force-independent factor $r_{u,\text{eff}}$ in the unbinding rate of the IF-microtubule bond is independent of the measuring geometry. The potential width $x_u$ enters the force-dependent factor of the unbinding rate and, thus, the force-sensitivity of the bond slightly increases from a vertical movement of the IF to a horizontal movement or a vertical movement with the microtubule turned by 45°.

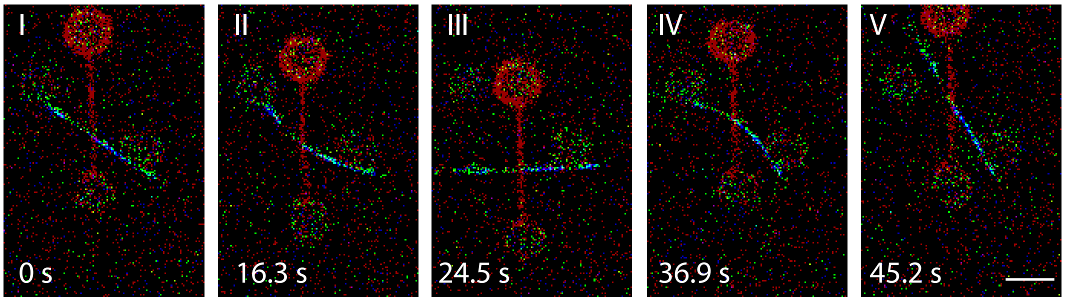


Figure 5: A strong IF-microtubule interaction that persisted for the downwards and upward pulling directions. The IF was first moved downward (I-II) until the microtubule broke off the right-hand-side bead (III). When the IF was moved upward, the microtubule re-attached to the right-hand-side bead (IV) and then broke off again (V), showing that the IF-microtubule interaction persisted even when pulling in different directions. Scale bar 5 µm.

### Movies

Movie 1: Confocal fluorescence movie of an IF-microtubule pair. We moved the IF vertically while it was in a perpendicular orientation with respect to the microtubule. The microtubule-IF pair interacted strongly; the microtubule broke off a bead. Scale bar 5 µm and 4.4 s per frame.

Movie 2: Confocal fluorescence movie of an IF-microtubule pair. We moved the IF at a 45°angle with respect to the microtubule. At first the filaments did not interact, then interactions became apparent by the deformation of the microtubule. At the end of the movie the microtubule broke off a bead due to strong interactions. Scale bar 5 µm and 2.4 s per frame.

Movie 3: Confocal fluorescence movie of an IF-microtubule pair. We moved the IF horizontally while it was in a perpendicular orientation with respect to the microtubule. The IF deformed while interacting with the microtubule. Eventually the interactions were so strong that the microtubule broke off the beads. Scale bar 5 µm and 2.2 s per frame.

Movie 4: Confocal fluorescence movie of an IF-microtubule pair. We moved the IF vertically while it was in a perpendicular orientation with respect to the microtubule. The IF-microtubule pair interacted, leading to deformations of the microtubule. Eventually the IF broke. Scale bar 5 µm and 5.6 s per frame.

Movie 5: Confocal fluorescence movie of an IF-microtubule pair. We moved the IF vertically while it was in a perpendicular orientation with respect to the microtubule. The

IF-microtubule did not interact. Scale bar 5 µm and 2.5 s per frame.

Movie 6 (same data as in Supplementary Figure 5): A strong IF-microtubule interaction that persisted for downwards and upward pulling direction. The IF was first moved downward until the microtubule broke off the right-hand-side bead. When the IF was moved upward, the microtubule re-attached to the right-hand-side bead and then broke off again, showing that the IF-microtubule interaction persisted even when pulling in different directions. Scale bar 5 µm and 8.4 s per frame.

### Modeling

#### Parameters for optical trapping experiments and modeling

Table 1: Parameters obtained from OT experiments (E), from modeling (M) and from literature (L).

| Parameter | Description | Value | E/M/L |
|---|---|---|---|
| $a_{\mathrm{IF}}$ | Diameter of a vimentin IF | 11 nm | L[1] |
| $a_{\mathrm{MT}}$ | Diameter of a microtubule | 25 nm | L[2] |
| $b(t)$ | Probability that an IF-microtubule bond is closed | | E, M |
| $\Delta G_{\mathrm{IF\text{-}MT}}$ | IF-microtubule bond energy in pure CB | $(2.3 \pm 1.3)\, k_B T$ | E, M |
| $\Delta G_{\mathrm{IF\text{-}MT,TX100}}$ | IF-microtubule bond energy with additional TX100 | $(2.1 \pm 0.6)\, k_B T$ | E, M |
| $\Delta G_{\mathrm{IF\text{-}MT,Mg}}$ | IF-microtubule bond energy with additional magnesium chloride | $(3.0 \pm 1.0)\, k_B T$ | E, M |
| $d_{\mathrm{IF}}$ | Length of the filament between the IF-microtubule interaction junction and bead 3 in OT experiments as sketched in Supplementary Fig. 3c | | E |
| $E_{Ab}$ | Binding activation energy of an IF-microtubule bond | | M |
| $E_{Au}$ | Unbinding activation energy of an IF-microtubule bond | | M |

| | | | |
|---|---|---|---|
| $f(t)$ | Density function of the exponential distribution to determine the time (un-)bindung of an IF-microtubule bond | | M |
| $F_B$ | Breaking force in OT experiments | | E |
| $F_e$ | Force increase during entropic stretching of a vimentin IF | | M |
| $k_B$ | Boltzmann constant | $1.38 \cdot 10^{-23}$ J/K | L[3] |
| $\lambda$ | Mean time until an (un-)binding event of an IF-microtubule bond | | M |
| $L_C$ | Contour length of a vimentin IF | | E |
| $L_P$ | Persistence length of a vimentin IF | 1.5 µm | L[4,5] |
| $l_{u,\mathrm{IF}}$ | Periodicity of vimentin filaments (length of a unit-length filament) | 43 nm | L[6] |
| $l_{u,\mathrm{MT}}$ | Periodicity of MTs (length of a tubulin dimer) | 8 nm | L[7] |
| $\nu$ | Probability of one interaction with a tubulin dimer within one persistence length of a vimentin IF | 1.1% | E |
| $n_i$ | Total number of IF-microtubule interactions of OT experiments in a specific buffer | | E |
| $n_{i,L_P}$ | Number of interaction sites within one persistence length of a vimentin IF | | E |
| $n_{\mathrm{pf}}$ | Number of protofilaments in a simulated microtubule | 13 | L[2] |

| | | | |
|---|---|---|---|
| $p_{\text{IF-MT}}$ | Probability of one microtubule subunit to interact with a vimentin IF subunit | | M |
| $p_b(t)$ | Probability that a bond closes at a certain time $t$ | | M |
| $p_u(t)$ | Probability that a bond opens at a certain time $t$ | | M |
| $p_{\text{ULF}}$ | Probability of a vimentin unit-length filament interacting with a tubulin dimer in an adjacent microtubule | | E |
| $r_{b,0}$ | Constant prefactor of the binding rate $r_b(t)$ | | M |
| $r_{b,\text{eff},y}$ | Effective binding rate in pure CB when the vimentin IF is moved vertically / in the $y$-direction and it is oriented perpendicularly to the microtubule | $1.07 \cdot 10^{-2}$ s$^{-1}$ | E |
| $r_{b,\text{eff},x}$ | Effective binding rate in pure CB when the vimentin IF is moved horizontally / in the $x$-direction and it is oriented perpendicularly to the microtubule | $2.4 \cdot 10^{-2}$ s$^{-1}$ | E |
| $r_{b,\text{eff},45^\circ}$ | Effective binding rate in pure CB when the vimentin IF is moved vertically / in the $y$-direction and it is oriented in a 45°angle to the microtubule | $1.6 \cdot 10^{-2}$ s$^{-1}$ | E |
| $r_{b,\text{eff},\text{TX100}}$ | Effective binding rate with additional TX100 | $0.56 \cdot 10^{-2}$ s$^{-1}$ | E |
| $r_{b,\text{eff},\text{Mg}}$ | Effective binding rate with additional magnesium chloride | $1.3 \cdot 10^{-2}$ s$^{-1}$ | E |

| | | | |
|---|---|---|---|
| $r_b(t)$ | Force-dependent binding rate of a vimentin IF and a microtubule | | M |
| $r_{u,0}$ | Constant prefactor of the unbinding rate $r_u(t)$ | | M |
| $r_{u,\text{eff},y}$ | Effective unbinding rate in pure CB when the vimentin IF is moved vertically / in the $y$-direction and it is oriented perpendicularly to the microtubule | $(0.11 \pm 0.10)$ s$^{-1}$ | M |
| $r_{u,\text{eff},x}$ | Effective unbinding rate in pure CB when the vimentin IF is moved horizontally / in the $x$-direction and it is oriented perpendicularly to the microtubule | $(0.12 \pm 0.09)$ s$^{-1}$ | E |
| $r_{u,\text{eff},45°}$ | Effective unbinding rate in pure CB when the vimentin IF is moved vertically / in the $y$-direction and it is oriented in a 45°angle to the microtubule | $(0.10 \pm 0.07)$ s$^{-1}$ | E |
| $r_{u,\text{eff,TX100}}$ | Effective unbinding rate with additional TX100 | $(0.26 \pm 0.20)$ s$^{-1}$ | M |
| $r_{u,\text{eff,Mg}}$ | Effective unbinding rate with additional magnesium chloride | $(0.15 \pm 0.08)$ s$^{-1}$ | M |
| $r_u(t)$ | Force-dependent unbinding rate of a microtubule and a vimentin IF | | M |
| $t$ | Time | | E, M |
| $t^*$ | Duration of entropic stretching of a vimentin IF in OT experiments | | M |

| | | | |
|---|---|---|---|
| $t_b$ | Time until formation of an IF-microtubule bond | | E, M |
| $t_{\mathrm{cont}}$ | Total time in which the IF and microtubule are unbound in OT experiments in a certain buffer | | E |
| $t_u$ | Duration of a closed IF-microtubule bond | | E, M |
| $\mathrm{d}t$ | Discretization time step | 0.05 s | M |
| $\tau$ | Time scale of force decrease in OT experiments | 0.1 s | E, M |
| $T$ | Temperature | | E, M |
| $v$ | Velocity of the lowest bead 3 as sketched in Supplementary Fig. 3 during OT experiments | 0.55 µm/s | E |
| $w$ | Final constant loading rate in OT experiments | | E |
| $x$ | End-to-end distance of a vimentin IF | | E, M |
| $x_u$ | Distance from bound to transition state in pure CB when the vimentin IF is moved vertically / in the $y$-direction and it is oriented perpendicularly to the microtubule | $(0.17 \pm 0.05)$ nm | M |
| $x_{u,x}$ | Distance from bound to transition state in pure CB when the vimentin IF is moved horizontally / in the $x$-direction and it is oriented perpendicularly to the microtubule | $(0.22 \pm 0.03)$ nm | M |

| | | | |
|---|---|---|---|
| $x_{u,45^\circ}$ | Distance from bound to transition state in pure CB when the vimentin IF is moved vertically / in the $y$-direction and it is oriented in a 45°angle to the microtubule | $(0.26 \pm 0.02)$ nm | M |
| $x_{u,\mathrm{TX100}}$ | Distance from bound to transition state with additional TX100 | $(0.23 \pm 0.13)$ nm | M |
| $x_{u,\mathrm{Mg}}$ | Distance from bound to transition state with additional magnesium chloride | $(0.08 \pm 0.08)$ nm | M |

#### Interaction probability of a tubulin and a vimentin subunit and geometry dependence

To compare the three binding rates of the three different measurement geometries in the OT experiments, we modeled the binding rates, which depend on the following parameters: The velocity $v$ with which periodic microtubule and vimentin subunits pass each other, the length $l_{u,\mathrm{IF}}$ of the unit-length filaments and the length $l_{u,\mathrm{MT}}$ of a tubulin dimer, the probability of a tubulin dimer and vimentin unit-length filament to bind to each other $p_{\text{IF-MT}}$, and the number of dimers and unit-length filaments in the overlapping area of both filaments $a_{\mathrm{IF}}/l_{u,\mathrm{MT}}$ or $a_{\mathrm{MT}}/l_{u,\mathrm{IF}}$. Thus, the binding rate for a vertical movement of the IF perpendicular to the microtubule is described as the product of an encounter rate and the probability $p_{\text{IF-MT}}$ that a bond is formed. The encounter rate in turn is given by the rate at which potential binding sites pass each other and the number of binding sites in the overlap area, which results in:

$$r_{b,\mathrm{eff},y} = \frac{v}{l_{u,\mathrm{IF}}} \frac{a_{\mathrm{IF}}}{l_{u,\mathrm{MT}}} p_{\text{IF-MT}}. \tag{1}$$

Note that here we describe binding between one subunit on each filament (tubulin dimer and vimentin unit-length filament, respectively). If binding involves contacts with several subunits on a filament, potential effective binding sites are bigger, but due to the periodic structure of the filaments there is the same number of binding sites per filament length and thus the same encounter rate. If there is more than one potential binding site within one filament subunit, the encounter rate is increased, and the inferred binding probability is reduced by the same factor, so that their product is the same. In all these cases, our binding parameters can be interpreted as effective parameters for the binding between a microtubule dimer and a vimentin unit-length filament. When the microtubule is turned by 45°, the binding rate is expected to change by a factor of $\sqrt{2}$ because the overlap area increases by a factor of $1/\cos(45°)$. Thus, the binding rate for the 45°-configuration with a vertical movement of the IF becomes:

$$r_{b,\mathrm{eff},45^\circ} = \frac{v}{l_{u,\mathrm{IF}}} \frac{a_{\mathrm{IF}}}{l_{u,\mathrm{MT}}} p_{\mathrm{IF\text{-}MT}} \sqrt{2}. \qquad (2)$$

In case of the horizontal movement of the IF along the microtubule, the rate of passing subunits changes to $v/l_{u,\mathrm{MT}}$ and the number of encounters of subunits is $a_{\mathrm{MT}}/l_{u,\mathrm{IF}}$:

$$r_{b,\mathrm{eff},x} = \frac{v}{l_{u,\mathrm{MT}}} \frac{a_{\mathrm{MT}}}{l_{u,\mathrm{IF}}} p_{\mathrm{IF\text{-}MT}}. \qquad (3)$$

We calculated the ratios of the binding rates for each pairing of measuring geometries. Comparison with the results from equations (1)-(3) shows good agreement:

$$\text{theoretical: } \frac{r_{b,\text{eff},y}}{r_{b,\text{eff},45^\circ}} = \frac{1}{\sqrt{2}} \simeq 0.71, \qquad \text{experimental: } 0.65$$

$$\text{theoretical: } \frac{r_{b,\text{eff},y}}{r_{b,\text{eff},x}} = \frac{a_{\text{IF}}}{a_{\text{MT}}} \simeq 0.44, \qquad \text{experimental: } 0.44,$$

$$\text{theoretical: } \frac{r_{b,\text{eff},45^\circ}}{r_{b,\text{eff},x}} = \frac{\sqrt{2}a_{\text{IF}}}{a_{\text{MT}}} \simeq 0.62, \qquad \text{experimental: } 0.67.$$

Alternatively, we can calculate the probability of a tubulin subunit to bind to a vimentin subunit from the experimental data using Eqs. (1)-(3):

$$\text{vertical movement, } 90^\circ\text{: } p_{\text{IF-MT}} \simeq 5.9 \cdot 10^{-4}$$

$$\text{vertical movement, } 45^\circ\text{: } p_{\text{IF-MT}} \simeq 6.4 \cdot 10^{-4}$$

$$\text{horizontal movement, } 90^\circ\text{: } p_{\text{IF-MT}} \simeq 6.0 \cdot 10^{-4}$$

The agreement of our geometric reasoning with the experimental results indicates that binding between microtubule and vimentin subunits does not depend on the time they stay in contact. By contrast, they bind rather rapidly once the subunits are in close proximity, but with low probability, suggesting that only a subset of subunits is able to bind. We hypothesize that controlling which subunits can bind (e.g. by posttranslational modifications) may provide a path for the cell to regulate the stabilization of microtubules by IFs.

#### Estimation of the bundling probability of microtubules and vimentin IFs

Our optical trapping experiments show a direct interaction between microtubules and vimentin intermediate filaments. However, we do not see co-alignment or bundling of the two filament types. To check whether such bundling should be expected, we estimate the probability of bundle formation between IFs and microtubules from the interaction probability of a vimentin unit-length filament with a tubulin dimer ($p_{\text{IF-MT}}$) determined in the previous section. Co-alignment requires interactions at more than one site within one persistence length of a vimentin filament to occur, since thermal fluctuations set the relevant length scale for tight contact between the filaments.

The probability $p_{\text{ULF}}$ of a vimentin unit-length filament interacting with a tubulin dimer in an adjacent microtubule is:

$$p_{\text{ULF}} = p_{\text{IF-MT}} \frac{l_{u,\text{IF}}}{l_{u,\text{MT}}} \frac{n_{\text{pf}}}{4} \simeq 0.011,$$

where $l_{u,\text{IF}} = 43$ nm is the length occupied by one vimentin unit-length filament within an IF, $l_{u,\text{MT}} = 8$ nm is the periodicity of tubulin dimers in the microtubule, $n_{\text{pf}} = 13$ is the number of protofilaments in a microtubule and $n_{\text{pf}}/4$ is the number of protofilaments per side facing the vimentin IF.

To obtain an upper limit for the probability of bundling, we assume that the two filaments that interact at one site are already aligned in parallel and estimate the probability of an additional interaction at a second site within one vimentin persistence length of the first site. This probability $\nu$ is obtained as

$$\nu \approx p_{\text{ULF}} \frac{L_P}{l_{u,\text{IF}}} \simeq 0.37\,. \tag{4}$$

with a persistence length of vimentin filaments of $L_P \simeq 1.5$ µm. If more than two interactions are required for bundling, the estimate is further decreased by $\nu^{(n_{i,L_P}-1)}$, for required interactions at $n_{i,L_P}$ sites. For $n_{i,L_P} = 2$, this leads to a probability of 14%. Thus, given the low probability of interaction, most microtubule-IF interactions would be mediated by a single site, even if the two filaments are pre-aligned. As filament pairs interacting at a single site can rotate relative to each other, the actual probabilities are even smaller. Thus, we would not expect a clear coalignment or bundling.

### Two-state model for IF-microtubule interactions

We modeled IF-microtubule interactions as single molecular bonds to understand the force-dependent behavior in different buffer conditions. The bond can either be in a closed or in an open state with force-dependent stochastic transitions between these two states, as sketched in Fig. 4b in the main text. In the experiment, we moved the IF with a constant speed $v$ perpendicularly to the microtubule, as shown in Fig. 2b in the main text and Supplementary Fig. 3a. Once the bond closed, the IF with an average persistence length of $L_P = 1.5$ µm[4,5] was stretched to its full contour length $L_C$. Thus, the entropic force $F_e$ relates to the end-to-end distance $x = vt$ as[8,9]

$$\frac{x}{d_{\mathrm{IF}}} = \coth\left(\frac{2L_P F_e}{k_B T}\right) - \frac{k_B T}{2L_P F_e} \tag{5}$$

with the Boltzmann constant $k_B$ and the temperature $T$. $d_{\mathrm{IF}}$ is the length of the filament between the IF-microtubule junction and bead 3 as sketched in Supplementary Fig. 3c.

In the simulation, we assumed a linear force increase from time $t^*$ on.[10,11] The linear force increase was set by the experimental force rate $w$, which we determined from a linear fit to the second half of the experimental force data of each interaction. $t^*$ was determined as the time when the force increase $\frac{dF_e}{dt}$ due to a decreasing entropy is the same as the experimental force rate $w$, i.e. $w = \frac{dF_e}{dt^*}$:

$$F(t) = \begin{cases} F_e(x = vt) & \text{for } \; t < t^* \\ wt & \text{for } t > t^* \end{cases} , \tag{6}$$

Once the bond broke at a force $F_B$ after a time $t_u$, we assumed an exponential force relaxation on a characteristic time scale $\tau$:

$$F(t) = F_B \exp(-(t - t_u)/\tau) \, \text{for } t > t_u. \tag{7}$$

We indeed observed a fast, exponential-like force decay in our experiments. However, the time resolution is not sufficient to fit $\tau$ precisely. We set $\tau = 0.1$ s as this results in force versus time curves similar to our experiments.

All variables with the index $b$ refer the binding process and the index $u$ represents the unbinding process. We describe the force-dependent binding and unbinding rates as follows: We assume that the binding and unbinding rates $r_b(t)$ and $r_u(t)$, respectively, depend on a reaction prefactor $r_{b,0/u,0}$, the activation energy for binding or unbinding $E_{Ab/Au}$, the thermal energy $k_BT$ and the potential width of the two states $x_{b/u}$:[12]

$$r_b(t) = r_{b,0} \exp\left(\frac{-E_{Ab}}{k_BT}\right) \cdot \exp\left(\frac{-F(t)x_b}{k_BT}\right), r_u(t) = r_{u,0} \exp\left(\frac{-E_{Au}}{k_BT}\right) \cdot \exp\left(\frac{F(t)x_u}{k_BT}\right) . \tag{8}$$

The force-independent parameters $r_{b,0/u,0}$ and $E_{Ab/Au}$ result in an effective zero-force rate

$$r_{b,\text{eff}/u,\text{eff}} = r_{b,0/u,0} \exp\left(\frac{-E_{Ab/Au}}{k_BT}\right) ,$$

in which $r_{b,\text{eff}}$ can be determined from the experimental data in Fig. 3c, e, g, i and k in the main text. We calculated the total contact time $t_{\text{cont}}$ of the IF and microtubule without an interaction and the number of initiated interactions $n_i$ between IFs and microtubules from the experimental data and get

$$r_{b,\mathrm{eff}} = \frac{n_i}{t_{\mathrm{cont}}} \, .$$

If we assume the same prefactor for the binding or unbinding process, i.e. $r_{b,0} = r_{u,0}$,[12] the ratio of two effective binding or unbinding rates for different experimental buffer conditions 1 and 2, or two different states (bound, unbound) sheds light on the differences in the activation energies for these buffer conditions or states. For the binding rates for two different buffer conditions, the activation energy difference is:

$$\frac{r_{b,\mathrm{eff},1}}{r_{b,\mathrm{eff},2}} = \frac{\exp\left(\frac{-E_{Ab,1}}{k_B T}\right)}{\exp\left(\frac{-E_{Ab,2}}{k_B T}\right)} \Rightarrow k_B T \ln\left(\frac{r_{b,\mathrm{eff},1}}{r_{b,\mathrm{eff},2}}\right) = E_{Ab,2} - E_{Ab,1} \, , \tag{9}$$

and likewise for the rates for the unbound state.

In the same way, we can calculate the absolute energy difference $\Delta G_{\text{IF-MT}}$ between the bound and unbound state for the same buffer condition:[12]

$$\Delta G_{\text{IF-MT}} = -k_B T \ln\left(\frac{r_{u,\mathrm{eff}}}{r_{b,\mathrm{eff}}}\right) \, . \tag{10}$$

Here, the sum of the potential widths $x_b + x_u$ nm provides the total distance between the bound and unbound state, which we assume to be the same for all experimental conditions. The rate equations in Eq. (8) ensure that detailed balance is satisfied.[12]

Thus, from these considerations and from the experiment, we know $L_C$, $w$, $\tau$ and $r_{b,\mathrm{eff}}$, but neither $r_{u,\mathrm{eff}}$ nor $x_b$ or $x_u$. We simulated the binding and unbinding reactions for the known parameters and varied $x_u$ from 0 nm up to 0.9 nm in steps of 0.01 nm and $r_{\mathrm{eff},u}$ from 0.02 up to 0.6 $\mathrm{s}^{-1}$ in steps of 0.01 $\mathrm{s}^{-1}$. We determined $x_b$ by calculating $x_b = 0.4\,\mathrm{nm} - x_u$, since the maximum value of $x_u$ is below 0.4 nm.

The binding and unbinding process cannot be described in a closed analytical expression due to the time dependence in the exponential expression of the reaction rates.[13] Therefore, we considered two different approaches to determine the breaking force histograms which we compared to the experimental data: (i) We solved the rate equations directly numerically, which is the fastest way to calculate the force histograms. (ii) We simulated the force-time trajectories of single bonds, which allows us to directly compare single simulated trajectories to our experimental data. Both approaches result in the same force histograms as shown in Fig. 3c, e, g, i and k in the main text, green shaded areas.

#### Numerical solution of the two-state Model

To solve the rate equations in Eq. (8) numerically, we defined $b(t)$ as the probability that the IF-microtubule bond is closed. Thus, the temporal behavior of $b$ can be described as:

$$\frac{\mathrm{d}b}{\mathrm{d}t} = -r_u(t) \cdot b(t) + r_b(t)(1 - b(t)) \,, \tag{11}$$

$$= -b(t) r_{u,\mathrm{eff}} \exp\left(\frac{F(t)x_u}{k_B T}\right) + (1 - b(t)) r_{b,\mathrm{eff}} \exp\left(\frac{-F(t)x_b}{k_B T}\right) . \tag{12}$$

We solved this expression numerically for $b(t)$ with the Matlab function `ode45`. To obtain a histogram of breaking forces, we differentiated $b(t)$ with respect to $t$ and, thus, determined the probability $p_u(t)$ that the IF and microtubule unbind at a certain time $t$:

$$p_u(t) = -\frac{\mathrm{d}b}{\mathrm{d}t} \,.$$

To calculate the probability-force diagram, to compare to the experiments, we determined $p_u$ as a function of $F$, i.e. $p_u(t(F))$ by inverting $F(t)$ as described in Eq. (6).

#### Monte-Carlo simulation of single molecular bonds described by the two-state model

To obtain single force-time trajectories of an IF-microtubule bond, we simulated the binding and unbinding process in several steps: (i) The time until an individual binding event was determined by choosing a random time $t_b$ from an exponential distribution with the density function $f(t)$ and a mean value of $\lambda = (r_b(t, F = 0))^{-1}$:[14,15]

$$f(t) = \lambda \exp(-\lambda t)\,.$$

The bond is now closed after time $t_b$. The force starts to increase as described in Eq. (6). (ii) As the unbinding rate depends on the force, which increases with time, the mean $(r_u(F(t)))^{-1}$ of the exponentially distributed unbinding time $t_u$ changes with increasing force. Thus, it is not straightforward to determine the time until unbinding with a single step as in (i). Instead, we split $t_u$ into small time intervals $\mathrm{d}t$. We set $\mathrm{d}t = 0.05$ s as a compromise between accuracy and computation time, which is the same as the experimental time resolution. The time was increased in steps of $\mathrm{d}t$ and after each step, the unbinding rate was evaluated. The probability $p_u$ that the bond breaks in the considered time interval is $p_u = r_u(F(t))\mathrm{d}t$, where we approximated the exponentially increasing unbinding rate as a constant for small $\mathrm{d}t$. If a random number drawn from a uniform distribution between 0 and 1 was greater than $p_u$, the bond stayed closed, otherwise it opened. If the bond remained closed, the time was increased by $\mathrm{d}t$, the force was updated and step (ii) was repeated until the bond broke. (iii) Once the bond broke, the force decreased as described by Eq. (7). Since the bond could close with a force-dependent rate while the force decayed, the time was increased stepwise again and the probability to rebind was evaluated as in step (ii) with $p_b = r_b(t, F(t))\mathrm{d}t$. If the force decreased to a value below 0.001 pN, the force was set to 0 pN and the algorithm was repeated starting at step (i).

As the IFs and microtubules had slightly different lengths for different measurements, the force rate differed between the experiments. To account for these different rates, we ran the simulation until 1000 breaking events are recorded for each experimental force rate $w$. The final distribution of breaking forces results from the normalized sum of distributions of breaking forces for all force rates. This final distribution was compared to the experimental data with the Kolmogorov-Smirnov test.[16] If the experimental and the simulated distributions did not differ more than allowed for a 5% significance level,[16] we accepted the parameters $r_{u,\mathrm{eff}}$ and $x_u$ as shown in Fig. 4a in the main text. To calculate the energy diagram in Fig. 4b in the main text, we determined the centroids of the accepted parameter regions in Fig. 4a in the main text. We determined the standard deviations from the distributions in Fig. 4a in the main text assuming that $r_{u,\mathrm{eff}}$ and $x_u$ are independent. The simulated breaking force histograms do not depend on the exact value of $x_b$ in the range of 0.2 to 1.5 nm since $r_{b,\mathrm{eff}}$ dominates over the force-dependent term in Eq. (8). We do not observe a sufficient number of rebinding events under force to determine $x_b$ from the experiment. For clarity, $x_b + x_u$ is set to 0.4 nm in Fig. 4 in the main text.

#### Parameters for TIRF experiments, modeling and simulations

Table 2: Parameters obtained from OT experiments (O), from TIRF experiments (T), from modeling of TIRF experiments (M), simulation input parameters (I) and parameters known from literature (L).

| Parameter | Description | Value | O/T/M/I/L |
|---|---|---|---|
| $c_{\mathrm{IF}}$ | Concentration of vimentin IFs in the TIRF experiment or the number of vimentin IFs per IF network volume | | E |
| $D$ | Average diffusion coefficient of a vimentin filament in the TIRF experiment | | L,[17] M |
| $\Delta G_{\mathrm{latd}}$ | Lateral association energy of a GDP dimer | 1.5 $k_BT$ | I |
| $\Delta G_{\mathrm{latt}}$ | Lateral association energy of a GTP dimer | 3.5 $k_BT$ | I |

| | | | |
|---|---|---|---|
| $\Delta G_{tb}$ | Total average energy of a tubulin dimer in the microtubule lattice before catastrophe at 20 µM tubulin and 2.3 µM vimentin | 7.1 $k_BT$ | O, T, M |
| $\Delta G_{tb}$ | Total average energy of a tubulin dimer in the microtubule lattice before catastrophe at 25 µM tubulin and 2.3 µM vimentin | 7.2 $k_BT$ | O, T, M |
| $\Delta G_{tb}$ | Total average energy of a tubulin dimer in the microtubule lattice before catastrophe at 20 µM tubulin and 3.6 µM vimentin | 5.7 $k_BT$ | O, T, M |
| $\Delta G_{tb}$ | Total average energy of a tubulin dimer in the microtubule lattice before catastrophe at 25 µM tubulin and 3.6 µM vimentin | 6.8 $k_BT$ | O, T, M |
| $f_{\text{cat, exp}}$ | Experimentally observed catastrophe frequency of microtubules with surrounding vimentin IFs at 20 µM tubulin and 2.3 µM vimentin | 0.123 $\text{min}^{-1}$ | T |
| $f_{\text{cat, exp}}$ | Experimentally observed catastrophe frequency of microtubules with surrounding vimentin IFs at 25 µM tubulin and 2.3 µM vimentin | 0.107 $\text{min}^{-1}$ | T |

| | | | |
|---|---|---|---|
| $f_{\text{cat, exp}}$ | Experimentally observed catastrophe frequency of microtubules with surrounding vimentin IFs at 20 µM tubulin and 3.6 µM vimentin | $0.111\ \text{min}^{-1}$ | T |
| $f_{\text{cat, exp}}$ | Experimentally observed catastrophe frequency of microtubules with surrounding vimentin IFs at 25 µM tubulin and 3.6 µM vimentin | $0.091\ \text{min}^{-1}$ | T |
| $f_{\text{cat, IF-MT}}$ | Catastrophe frequency of microtubules while interacting with a vimentin IF | | M |
| $f_{\text{cat, MT}}$ | Experimentally observed catastrophe frequency of microtubules | $0.180\ \text{min}^{-1}$ (20 µM), 0.156 $\text{min}^{-1}$ (25 µM) | T |
| $f_{\text{resc}}$ | Simulation rescue frequency at 25 µM tubulin without IFs | $0.03\ \text{s}^{-1}$ | I |
| $f_{\text{resc, IF}}$ | Simulation rescue frequency at 25 µM tubulin with IFs at a concentration of 2.3 µM | $0.17\ \text{s}^{-1}$ | I |
| $\zeta$ | Mesh size of the vimentin network in TIRF experiments | 0.63 µm (2.3 µM vimentin), 0.55 µm (3.6 µM vimentin) | E, L[18,19] |
| $\eta$ | Viscosity of sample studied in TIRF experiments | 3 mPas | L[20] |
| $M$ | Number of tubulin dimers which are bound to a vimentin filament subunit at the same time | | M |

| | | | |
|---|---|---|---|
| $n$ | Number of lateral neighbors of a tubulin dimer | | I |
| $n_{\mathrm{pf}}$ | Number of protofilaments in a simulated microtubule | 13 | L[2] |
| $p_i$ | Probability that a vimentin monomer interacts with a tubulin dimer | 33% | O, T, M |
| $r$ | Random number between 0 and 1 | | I |
| $r_{\mathrm{diff}}$ | Diffusion limited encounter rate of vimentin IFs and microtubules | $90\ \mathrm{s}^{-1}$ | M |
| $r_{dd,0}$ | Depolymerization rate of a GDP dimer without lateral neighbors | $643\ \mathrm{s}^{-1}$ | I |
| $r_{dt,0}$ | Depolymerization rate of a GTP dimer without lateral neighbors | $9.93 \cdot 10^{-4}\ \mathrm{s}^{-1}$ | I |
| $r_{dd}$ | Depolymerization rate of a GDP dimer taking the number of neighbor dimers into account | | I |
| $r_{dt}$ | Depolymerization rate of a GTP dimer taking the number of neighbor dimers into account | | I |
| $r_{hy}$ | Hydrolysis rate of GTP dimers | $7\ \mathrm{s}^{-1}$ | I |
| $r_i$ | Interaction rate of IFs and microtubules in the TIRF experiments | $0.06\ \mathrm{s}^{-1}$ | M |
| $r_{g,20}$ | Polymerization rate of GTP dimers per protofilament for 20 µM free tubulin concentration | $1.3\ \mathrm{s}^{-1}$ | I |

| | | | |
|---|---|---|---|
| $r_{g,25}$ | Polymerization rate of GTP dimers per protofilament for 25 µM free tubulin concentration | 2.2 $s^{-1}$ | I |
| R | Any reaction rate in simulation | | I |
| $z$ | Random number between 0 and 1 | | I |

#### Model of a dynamic microtubule

We based our model of a dynamic microtubule on Refs. 21 and 22 and ran Monte-Carlo simulations with a self-written Python code (Beaverton, OR, USA) to obtain simulated kymographs. We assumed a microtubule lattice with $n_{\mathrm{pf}} = 13$ protofilaments that has a helical pitch of 3 monomers per turn as sketched in Fig. 5a in the main text. Thus, there is a seam formed by protofilaments 1 and 13, which are displaced by 1.5 dimers. All dimers incorporated in the lattice interact with two lateral and two longitudinal dimer positions. At the seam, the dimers interact with two half dimers across the seam. The microtubule is represented by a matrix in the simulation and the state of the dimer is entered at a corresponding position in the matrix. A dimer position can be either unoccupied or occupied by a GTP dimer (purple in Fig. 5a, b in the main text), a GDP dimer (blue in Fig. 5a, b in the main text) or a GMPCPP-dimer (green in Fig. 5a, b in the main text). We set the first three dimer layers to GMPCPP dimers, which represent the seed in the experiment. The GMPCPP dimers cannot depolymerize. To avoid artifacts from the starting conditions, we started the simulations with a microtubule consisting of 30 layers of GDP dimers, which have four layers of GTP dimers on top representing the tip.[21]

To simulate microtubule dynamics, we determined four different reaction rates (i–iv) as sketched in Fig. 5a (top) in the main text: (i) The polymerization rate $r_g$ when a GTP dimer binds to the tip of the microtubule, (ii) the depolymerization rate $r_{dt}$ of a GTP dimer when a GTP dimer falls off the lattice, (iii) the hydrolysis rate $r_{hy}$ of a GTP dimer to a GDP dimer and (iv) the depolymerization rate $r_{dd}$ of GDP dimers. Since we used a buffer which is also compatible with vimentin filament assembly, these simulation parameters differ from the parameters used in literature.[21,23,24] We summarize all important simulation parameters in Table 2. We calculated the different reaction rates (i–iv) as follows:

(i) The polymerization rate for GTP dimers is concentration dependent.[21] To match the growth rate to the experimentally observed one, we set it to $r_{g,20} = 1.3$ dimers $s^{-1}$ per protofilament for 20 µM free tubulin concentration and to $r_{g,25} = 2.2$ dimers $s^{-1}$ per protofilament for 25 µM free tubulin concentration.

(ii)/(iv) The depolymerization rate of GTP and GDP dimers depends on the number of lateral neighbors $n$. For each lateral dimer, the depolymerization rate was lowered by a factor of $\exp(-\Delta G_{\text{latt/latd}})$ due to the change in total bond energy $\Delta G_{\text{latt}} = 3.5\ k_BT$ for a GTP dimer and $\Delta G_{\text{latd}} = 1.5\ k_BT$ for a GDP dimer:[21]

$$r_{dt/dd} = r_{dt/dd,0} \exp\left(\frac{-n\Delta G_{\text{latt/latd}}}{k_BT}\right) , \tag{13}$$

For no lateral dimers, we assumed an unbinding rate of $r_{dt,0} = 9.93 \cdot 10^{-4}\ s^{-1}$ for GTP and $r_{dd,0} = 643\ s^{-1}$ for GDP. We assumed that only the dimers at the tip of a protofilament can depolymerize.[22]

(iii) We set the hydrolysis rate to $7\ s^{-1}$ to obtain a tip size which results in the same change in catastrophe frequency as observed in our experiments. This rate is on the same order of magnitude as assumed in Ref. 21. A dimer can only hydrolyze, if it has a neighbor in the same protofilament towards the direction of growth.[21,22] Since we did not observed rescue in our experiments for a free tubulin concentration of 20 µM and the precise reason for rescue is unknown,[23] we assumed that the rapidly disassembling microtubule is "locked" in the disassembly state and no rescue occurs because GTP dimers polymerize faster then GDP dimers depolymerize.[23] Yet, we observe rescue at a concentration of 25 µM, which we implemented in our simulation as occurring with a rate of $f_{\text{resc}} = 0.03\ s^{-1}$.[23]

To simulate a kymograph of a dynamic microtubule, we calculated all possible reaction rates. For each possible reaction with rate $R$, a random number $z$ between 0 and 1 was drawn, with which we determined the time until the next realization of a certain reaction:[14,21]

$$t = \frac{-\ln z}{R}\,. \tag{14}$$

The reaction with the smallest time was set to be the next occurring reaction. The microtubule matrix containing the dimer states was updated correspondingly as shown for a snapshot of a typical microtubule configuration in Fig. 5b in the main text. We ran 100 simulations for a total simulated time of 900 s each to obtain comparable amounts of experimental and simulated data. We recorded the length of the shortest protofilament during the simulation, which results in simulated kymographs. We plotted typical simulated kymographs in Fig. 5c (left) in the main text for 20 µM free tubulin without surrounding vimentin and in Fig. 5c (right) in the main text for 25 µM free tubulin with surrounding vimentin.

#### Model of a dynamic microtubule stabilized by IFs

The above model of a dynamic microtubule was modified in the following way to account for the direct binding of IFs as seen in our OT experiments: We assumed that IFs bind stochastically to the microtubule lattice. We note that from our experiments, we cannot make precise conclusions about the molecular mechanism causing the interaction, therefore the molecular mechanism is not specified in our model. We hypothesize that IFs bind to individual tubulin dimers, but based on our experiments we cannot exclude the possibility that the interaction is based on larger binding sites that consist of multiple tubulin dimers. However, in the OT experiments, we always observed that the bond between an IF and a microtubules broke in a single step, so that if binding involves multiple tubulin dimers, it must be highly cooperative and can still be treated effectively as a single bond. The rates of binding and unbinding of IFs to the microtubule were calculated from those determined in the OT experiments, accounting for the different geometry in the TIRF approach. This calculation is described at the end of this section. We further assumed that the presence of a bound IF modulates the depolymerization rates $r_{dt/dd}$, but does not affect the polymerization and hydrolysis rates.

*Depolymerization rates:* Our OT experiments showed that IFs directly interact with microtubules. By comparing the binding and unbinding rates (the latter in the force-free limit), we determined the energy difference $\Delta G_{\text{IF-MT}}$ between the bound and unbound state of the IF-microtubule interactions. Thus, if an IF binds to a microtubule dimer, the total binding energy of the dimer in the microtubule lattice is increased by $\Delta G_{\text{IF-MT}}$, which lowers the total energy sum in the exponential term of Eq. (13),

$$r_{dt/dd} = r_{dt,0/dd,0} \exp\left(\frac{-n\Delta G_{\text{latt/latd}} - \Delta G_{\text{IF-MT}}}{k_B T}\right) \tag{15}$$

and thus reduces the depolymerization rate, specifically for the case of GTP-dimers in the microtubule cap, where the depolymerization rate is small anyway. This assumption can be interpreted as follows: when a tubulin dimer to which an IF is bound unbinds from the microtubule lattice, the IF also unbinds. Based on our experiments, we cannot distinguish whether the IF is bound to a single tubulin dimer or to multiple dimers, but we know that if the latter case applies, unbinding from those dimers must be cooperative since we do not observe step-wise unbinding in OT experiments. Therefore, the same model applies to both scenarios, i.e., if IF-microtubule binding involves more than one dimer, unbinding of one of those dimers also unbinds the IF from the other dimers. The only difference between the scenarios is that the binding energy per dimer is $\Delta G_{\text{IF-MT}}/M$ if $M$ dimers contribute to the bond. Due to the cooperativity, however, the total energy $\Delta G_{\text{IF-MT}}$ enters the depolymerization rate.

*Binding rate of vimentin IFs to microtubules:* From the OT experiments, we know that the binding rates of IFs and microtubules depend on the geometry of the experiment, i.e. the IF is moved perpendicularly vertically or horizontally compared to the microtubule or at an angle. To obtain the binding rate in the geometry of the TIRF experiments, we used the same approach as in the section "Interaction probability of a tubulin and a vimentin subunit" describing the binding rate by a geometry-dependent encounter rate and a geometry-independent binding probability. In the TIRF experiments, the encounter rate is different compared to the OT experiments since the vimentin filaments diffuse and are not moved in a certain direction relative to the microtubule. Therefore, we calculated the diffusion limited encounter rate[25] $r_{\text{diff}}$ of vimentin and microtubule subunits:

First, we determined the average diffusion coefficient of the vimentin IFs: The estimated viscosity[26] $\eta \simeq 3$ mPas of the sample in TIRF experiments deviates from the viscosity of water, since the sample in TIRF experiments contained 0.09% methylcellulose. The diameter[1] $a_{\text{IF}}$ of a vimentin filament is 11 nm. The diffusion occurs in three dimensions with a diffusion coefficient[17] of $D = k_B T \ln(\zeta/a_{\text{IF}})/(3\pi\zeta\eta)$.

Second, we estimated the concentration $c_{\text{IF}}$ of vimentin IFs in the network or the number of vimentin IFs per network volume as $c_{\text{IF}} = 3\zeta/a_{\text{IF}}/\zeta^3$, where $\zeta \simeq 0.63$ µm or $\zeta \simeq 0.5$ µm is the mesh size of the vimentin filament network[18] for 2.3 µM or 3.6 µM vimentin, respectively.

Third, we calculated the diffusion limited encounter rate,[25] taking into account the diameter of a vimentin IF $a_{\text{IF}} \simeq 11$ nm:[1]

$$\begin{aligned} r_{\text{diff}} = 4\pi D a_{\text{IF}} c_{\text{IF}} &\simeq 90\,\text{s}^{-1} \text{ for } 2.3\text{ µM vimentin}\,, \\ &\simeq 150\,\text{s}^{-1} \text{ for } 3.6\text{ µM vimentin}\,. \end{aligned}$$

To determine the interaction rate $r_i$ of a microtubule subunit and a vimentin filament subunit, we multiplied the encounter rate $r_{\text{diff}}$ with the interaction probability of a microtubule subunit and a vimentin filament $p_{\text{IF-MT}}$ that was already determined from the OT experiments :

$$r_i = r_{\text{diff}}\, p_{\text{IF-MT}} \simeq 0.06\,\text{s}^{-1} \text{ for } 2.3\ \mu\text{M vimentin}\,,$$
$$\simeq 0.09\,\text{s}^{-1} \text{ for } 3.6\ \mu\text{M vimentin}\,.$$

We calculated the probability $p_i$ that an IF is bound to a microtubule by assuming an equilibrium between binding and unbinding IFs:

$$r_i(1-p_i) = r_{u,\text{eff}} p_i\,.$$

The unbinding rate was determined from OT experiments as well. We find $p_i \simeq 33\%$ in case of 2.3 µM vimentin and $p_i \simeq 44\ \%$ in case of 3.6 µM vimentin. Consequently, in our simulation, we drew a random number $r$ between 0 and 1 and if $r < p_i$, the depolymerization rate changes as described in Eq. (15). If $r > p_i$, the depolymerization rate remains unchanged.

The additional binding energy of IFs to microtubules also decreases the depolymerization rate of potential rescue sites, thus, rescue occurs more often. Thus, the frequency for rescue sites with surrounding vimentin filaments increases from $f_{\text{resc}} = 0.03\ \text{s}^{-1}$ to $f_{\text{resc, IF}} = 0.17\ \text{s}^{-1}$ in case of 2.3 µM vimentin. The rescue frequency of microtubules with surrounding filaments is lower than we would expect if we calculate $f_{\text{resc}} \exp(\Delta G_{\text{IF-MT}}/k_B T)\ \text{s}^{-1} = 0.3\ \text{s}^{-1}$, however, on the same order of magnitude. Our model is probably too simple to describe this discrepancy arising from the poorly understood rescue process.[23]

#### Estimate of tubulin dimer binding energy by combining results from optical trapping and TIRF experiments

We can estimate the tubulin dimer binding energy by combining the results from OT and TIRF experiments. First, we calculated the catastrophe frequency $f_{\text{cat, IF-MT}}$ of a microtubule when a vimentin filament continuously interacts with all dimers. We know the experimentally observed catastrophe frequency without vimentin in solution $f_{\text{cat,MT}}$ and with vimentin in solution $f_{\text{cat,exp}}$ from the TIRF experiments. The observed catastrophe frequency in presence of vimentin results from a combination of microtubules which are in contact with a vimentin IF and microtubules which are not in contact with an IF. The probability that a microtubule monomer and a vimentin IF are in contact is $p_i$. Thus, the observed catastrophe rate $f_{\text{cat,exp}}$ in presence of vimentin IFs and the catastrophe rate $f_{\text{cat, IF-MT}}$ for microtubules continuously interacting with a vimentin IF are:

$$\begin{aligned}
f_{\text{cat,exp}} &= (1-p_i) f_{\text{cat, MT}} + p_i f_{\text{cat, IF-MT}} \\
f_{\text{cat, IF-MT}} &= \frac{f_{\text{cat,exp}} - (1-p_i) f_{\text{cat, MT}}}{p_i} \\
&\simeq 0.0056\,\text{min}^{-1} \text{ for } 20\text{ µM and } 0.0061\text{ min}^{-1} \text{ for } 25\text{ µM tubulin, } 2.3\text{ µM vimentin,} \\
&\simeq 0.022\,\text{min}^{-1} \text{ for } 20\text{ µM and } 0.0073\text{ min}^{-1} \text{ for } 25\text{ µM tubulin, } 3.6\text{ µM vimentin.}
\end{aligned}$$

During depolymerization of the microtubule, the additional energy of a GTP dimer in the microtubule lattice $\Delta G_{tb}$ is released. Therefore, we assumed that the only energy difference between the dimer, which is incorporated in an microtubule and which unbinds from an IF monomer in the OT experiments, and the last dimer, which depolymerizes just before an microtubule catastrophe in the TIRF experiments, is $\Delta G_{tb}$. Thus, we can combine the catastrophe rates from TIRF experiments and the unbinding rates of the OT experiments to calculate $\Delta G_{tb}$:

$$
\begin{aligned}
\frac{r_{u,\text{eff}}}{f_{\text{cat,IF-MT}}} &= \exp\left(\frac{\Delta G_{tb}}{k_B T}\right) , \qquad (16)\\
\Delta G_{tb} &= k_B T \ln\left(\frac{r_{u,\text{eff}}}{f_{\text{cat,IF-MT}}}\right)\\
&\simeq 7.1\, k_B T \text{ for } 20\,\mu\text{M and } 7.2\, k_B T \text{ for } 25\,\mu\text{M, } 2.3\ \mu\text{M vimentin,}\\
&\simeq 5.7\, k_B T \text{ for } 20\,\mu\text{M and } 6.8\, k_B T \text{ for } 25\,\mu\text{M, } 3.6\ \mu\text{M vimentin.}
\end{aligned}
$$

### Code for simulations

#### Simulation of single intermediate filament-microtubule interactions

```
 % simulate two crossed and binding filaments

% entropic stretching at beginning included which depends on the
    length of
% the filaments involved

close all
clear all
clc

% ----------------------------------------------------------------
%% 1. set parameters
% ----------------------------------------------------------------

% load force rates and lengths of filament lengths
fileID=fopen('0Mg0Tx_fillengths.txt','r');
l_set=fscanf(fileID, '%f\n')';
fclose(fileID);

fileID=fopen('0Mg0Tx_lr.txt','r');
v_set=abs(fscanf(fileID, '%f\n')');
fclose(fileID);

% set moving speed of bead pair 3+4
v_bp=0.55; % um/s

% thermal energy
kbt=4.1; % in units of kbt = 4.1 pN nm

% effective closing rate from experiment
r_B0= 0.0107; % in 1/s

% approximate effective unbinding rate from experiment
r_U0_set=0.03:0.01:0.06; % in 1/s
```

```
% effective potentiel width during unbinding
x_Ueff_set=0.5:0.01:0.57; % in nm

% effective potential width during binding
x_Beff_set=0.4-x_Ueff_set; % in nm

% force decay factor for exponential force loss when unbound
tau=0.01; % in s

% estimate time between closing and breaking of a bond
dt=1/20; % compromise between accuracy and experimental resolution

% maximum possible force to estimate for which forces the
    unbinding rate
% needs to be calculated
F_max=7000;

% maximum time in s of how long a bond stays closed
t_max=F_max/20; % s

% number of time intervals
t_nr=t_max/dt;

% time intervals
t=linspace(0,t_max, t_nr);

l_p=1.5; % um, persistence lenght

%------------------------------------------------------------
%% 2. simulate force-time curves
%------------------------------------------------------------
% run a loop for the number of experiments carried out; each
    experiment
% results in a force-time curve

for r_U0=r_U0_set
for x_Ueff=x_Ueff_set
    x_Beff=1-x_Ueff;

    % number of repetitions of force time trajectory recording
    n_exp_total=200;

    % number of conditin:
    nr_l=0;

    foldername=['xu_',num2str(x_Ueff), '_ruEff_', num2str(r_U0)];
    mkdir(['approximate_simulation_to_experiment_vimMT/0Mg0Tx/...
    param_sweep_high_xu_small_ru/', foldername])

for v=v_set(1:end)
    %% calculate force rate lookup table
    % clear all variables from the old lengths and loading rate
    clear exten_entr t_entr t_entr_dt f_entr_interp
       exten_entr_interp F_lu
    nr_l=nr_l+1; % new lengths/force rates have to be loaded
    % create force vector
    force_entr=linspace(0, 5, 100); % pN
```

```
% calculate extension
exten_entr=l_set(nr_l).*(coth(2.*l_p.*force_entr.*10^3./(kbt))
    - ...
kbt./(2.*l_p.*force_entr.*10^3));
% delete all NaN values as preparation for interpolation
force_entr(isnan(exten_entr))=[];
exten_entr(isnan(exten_entr))=[];
% interpolate time and force to distances in dt for the time
t_entr=(exten_entr-exten_entr(1))/v_bp; % time in s
t_entr_dt=linspace(0, max(t_entr), max(t_entr)/dt);
% make equally distanced time vector for interpolation
f_entr_interp=interp1(t_entr, force_entr, t_entr_dt);
exten_entr_interp=interp1(t_entr, exten_entr, t_entr_dt);

% start when force has reached 0.01 pN and set all parameters
    to start at
% this value so that if the junction is bound, it does not
    take ages to
% increase the force
idx_minF=find(min(abs(f_entr_interp -0.01))==abs(f_entr_interp
    -0.01));
t_entr_dt=t_entr_dt(idx_minF:end);
f_entr_interp=f_entr_interp(idx_minF:end);
exten_entr_interp=exten_entr_interp(idx_minF:end);

% find force rate for the entropic stretching and match it to
    the linear
% rate
frate_eff=(f_entr_interp(2:end)-f_entr_interp(1:end-1)). ...
/(t_entr_dt(2:end)-t_entr_dt(1:end-1));

% if the entropic stretching part is very small, we assume
    that the
% slope directly linearly increases
if isempty(frate_eff)
    F_lu=v.*t;
else
    % find force rate which is the same as in the experiment
    idx_entr_end=min(find(min(abs(frate_eff-v))==abs(frate_eff
        -v)));
    % min-function of minima: to find earliest minimum
    % after that index, start the linear force increase //
        F_lu=F look up
    F_lu=[f_entr_interp(1:idx_entr_end), v.*t(idx_entr_end+1:
        end)- ...
    v.*t(idx_entr_end)+f_entr_interp(idx_entr_end)];
end

tic
% determine opening rate
for k=1:t_nr
    r_o(k)=exp(x_Ueff.*F_lu(k)./kbt).*r_U0;
end
```

```
% variable for saving force jumps
f_jump_all=[];

%% start actual run of experiment
for n_exp=1:n_exp_total
time=0; % set starting time to be 0
time_f=0; % time_f is the duration for which the force is > 0
steps=1; % number of steps of the Gillespie algorithm
clear F RATES OPENS CLOSES delta_t Nt timet state f_jump
   counts
% state of the MT / IF junction
state=0; % state of the bond; ...
% 0: unbound/open/no interaction, 1: bound/closed/interaction
F=0; % force is 0 at beginning
F_hr=0; % "high resolution" force which includes the
   exponential
% force decay
t_hr=0; % "high resolution" time which includes exponential
   force decay
s_hr=0; % "high resolution" state of the bond

% count number of force jumps
counter_fj=1;
while counter_fj<6 % how many force jumps we want to record
     % check whether the bond is closed or open
    if state(steps)==0 % bond is open

        % progress the reaction time
        delta_t(steps)=exprnd(1/r_B0);

        % increase the number of Gillespie steps
        steps=steps+1;

        % increase the time by the time step delta_t
        time(steps)=time(steps-1)+delta_t(steps-1);

        % set force
        F(steps)=0;

        % change the state of the bond to bound; this happens
           after
        % increasing the number of Gillespie steps to make
           sure the
        % bond is closed at first: state(1)=0, state(2)=1;
        state(steps)=1;

        % record force, time and state
        F_hr=[F_hr, F(steps)];
        t_hr=[t_hr,time(steps)];
        s_hr=[s_hr, state(steps)];
    else % bond is closed
        % introduce a counter for the steps to make until the
           bond is
        % open; "help step counter"
        steps_h=1;
```

```
% introduce also a helping variable for saving the
    state
% during the small time steps dt
state_h=1; % bond is closed

% ———————————
% WHILE LOOP FOR FORCE INCREASE
% ———————————
while state_h(steps_h)==1 % increase force only when
    bond is closed
    % calculate p_o, the probability that the bond
        will
    % open in a certain time interval dt
    p_o=r_o(steps_h+1).*dt;

    % draw a number from an equal distribution between
        0
    % and 1 to see whether the reaction takes place
    rdm=rand;

    if rdm<p_o % bond opens
        state_h(steps_h)=0; % bond open
        steps=steps+1; % increase step in overall
            simulation
        time(steps)=time(steps-1)+steps_h*dt; % add
            time to
        % entire force-time curve
        state(steps)=0; % change state of the system
            to open
        F(steps)=F_lu(steps_h);%F(steps-1)+v.*steps_h
            .*dt;

        % save system force, time, state
        F_hr=[F_hr, F(steps)];
        t_hr=[t_hr, time(steps)];
        s_hr=[s_hr, state(steps)];
        f_jump(counter_fj)=F(steps);
        counter_fj=counter_fj+1;
    else % if the bond stays closed
        F_hr=[F_hr, F_lu(steps_h)];
        t_hr=[t_hr, t_hr(end)+dt];
        steps_h=steps_h+1; % increase step to make
            another
        % small dt step
        state_h(steps_h)=1; % bond stays closed
    end
end

% ——————
% WHILE LOOP FOR FORCE DECREASE
% ——————
% initialize helpful variables
steps_h=1;
```

```
F_h=F(steps);
time_h=0;
state_h=0;
while state_h(steps_h)==0 % force relaxation only
% when bond is open
    % calculate force-depending closing rate from
       current force
    r_c= r_B0.*exp(x_Beff*F_h(steps_h)/kbt);

    % calculate p_c, the probability that the bond
       will close in a
    % certain time interval dt
    p_c=r_c*dt;

    % draw a number from an equal distribution between
       0 and 1
    % to see whether the reaction takes place
    rdm=rand;
    if rdm<p_c % bond closes
        steps_h=steps_h+1;
        state_h(steps_h)=1; % bond closed
        steps=steps+1; % increase step in overall
           simulation
        time(steps)=time(steps-1)+steps_h*dt; % add
           time to entire force-time curve
        state(steps)=1; % change state of the system
           to close
        F(steps)=F(steps-1).*exp(-steps_h*dt/tau) ;

        % save system force, time, state
        F_hr=[F_hr, F(steps)];
        t_hr=[t_hr, time(steps)];
        s_hr=[s_hr, state_h(steps_h)];
    else % bond stays open
        steps_h=steps_h+1; % prepare to make another
           small dt step
        state_h(steps_h)=0; % bond stays open
        F_h(steps_h)=F_h(steps_h-1).*exp(-dt/tau);

        % save system force, time, state
        F_hr=[F_hr, F_h(steps_h)];
        t_hr=[t_hr, t_hr(end)+dt];
        s_hr=[s_hr, state_h(steps_h)];
    end

    if state_h(steps_h)==0 && F_h(steps_h)<0.001 %
       force is so small that we can set it to 0 and
       continue with normal Gillespie
        steps=steps+1;
        state(steps)=0;
        F(steps)=0;

        % save system force, time, state
        F_hr=[F_hr, F(steps)];
```

```
                time(steps)=time(steps-1)+steps_h*dt;
                t_hr=[t_hr, time(steps)];
                s_hr=[s_hr, state(steps)];
                break % break the while loop for the
                    decreasing force and
                % continue for-loop and determine the time
                    when the bond closes with one large
                    Gillespie step
            end
        end
    end
end

f_jump_all=[f_jump_all, f_jump];
end

%% plot typical data set
figure
plot(t_hr,F_hr, '.')
print(['approximate_simulation_to_experiment_vimMT/0Mg0Tx/
    param_sweep_high_xu_small_ru/',foldername,'/typicalF_v',
    num2str(v), '_rB0', ...
     num2str(r_B0), '_rU0', num2str(r_U0), '_xBeff',num2str(
        x_Beff), '_xUeff', num2str(x_Ueff)],'-dpng', '-r400')

%% calculate histogram of force jumps

F_max_jump=max(f_jump_all);

% plot histogram for force jumps
edges=linspace(0, 500, 500/5); % bin size: 5 pN, maximum force
    : 100 pN
forces=linspace((edges(2)-edges(1))/2,(edges(end-1)+edges(end)
    )/2, length(edges)-1);
figure
hist=histogram(f_jump_all, edges);
counts=hist.BinCounts;
close all

figure
hold on
x=bar(forces, counts/sum(counts), 'DisplayName', ['v=',num2str
    (v),', r_{B0}=', ...
     num2str(r_B0), ', r_{U0}=', num2str(r_U0),', x_{Beff}=',
        num2str(x_Beff),', x_{Ueff}=',...
     num2str(x_Ueff)])
legend('Location', 'Northeast')
ylabel('relative occurence')
xlabel('force (arb. units)')
box on
title('all breaking forces')
fig = gcf;
fig.PaperUnits = 'inches';
fig.PaperPosition = [0 0 7 3];
```

```
    print(['approximate_simulation_to_experiment_vimMT/0Mg0Tx/
       param_sweep_high_xu_small_ru/',foldername,'/v', num2str(v),
        '_rB0', ...
        num2str(r_B0), '_rU0', num2str(r_U0), '_xBeff',num2str(
           x_Beff), '_xUeff', num2str(x_Ueff)],'-dpng', '-r400')

    %% save data
    close all
    save(['approximate_simulation_to_experiment_vimMT/0Mg0Tx/
       param_sweep_high_xu_small_ru/',foldername,'/v', num2str(v),
        '_rB0', ...
    num2str(r_B0), '_rU0', num2str(r_U0), '_xBeff',num2str(x_Beff)
       , '_xUeff', num2str(x_Ueff),'.mat'])
    toc

end
end
end
```

#### Simulation of a dynamic microtubule interacting with a vimentin intermediate filament

```
#!/usr/bin/env python3
# -*- coding: utf-8 -*-
"""
Created on Thu Dec  5 17:16:31 2019

@author: clorenz
"""
import numpy
import matplotlib.pyplot as plt # for plotting
import random # for drawing random numbers
import os # for making new folders
import time # to measure time
import sys #stop system running

# name for folder to save files in
savename='test/'

# ---------------------------------------------------------------------------
# parameter for number of simulation runs and basic MT
   architecture
# ---------------------------------------------------------------------------

# define parameters for simulation
nr_pf=13 # total number of protofilaments

# maximum length of microtubule, artificially set for this
   simulation
max_mt_l=2500

# length of seed
nr_seed=4
```

```
# length of GTP tip at beginning
lgtptip=4

# length of GDP shaft at beginning
lgdpshaft=30

# experimental parameter: tubulin concentration
c_tb=25 # uM

# ------------------------------------------------------------------------
# reaction rates
# ------------------------------------------------------------------------
# hydrolization rate
k_hy=7 # 1/s


# on rate for GTP dimer; addition is only possible
# when there is a longitudinal dimer
k_on_base=2.2 # 1/s
k_on=[k_on_base, k_on_base, k_on_base]#37*c_tb, 25*c_tb, 1.7*c_tb]
    # 1/s; for 0 lateral neighbors, 1 lateral neighbor
# and 2 lateral neighbors in this order

# k_on at the seem:
k_on_seem=[k_on_base, k_on_base, k_on_base, k_on_base] # for 0
   lateral dimers, 1/2 dimer, 1 dimer, 1.5 laterl dimers and 2
   lateral dimers


# off rates for different numbers of lateral neighbors for GTP
dG_latgtp=3.5 # lateral bond energy per bond per dimer
# ... no lateral neighbor:
k_offgtpbase=0.00003 # basic rate for all dimers 1/s
k_offgtp=[k_offgtpbase*numpy.exp(dG_latgtp), k_offgtpbase,
   k_offgtpbase*numpy.exp(-dG_latgtp)]#[3500,25, 10] # 1/s; for 0
   lateral neighbors, 1 lateral neighbor
# and 2 lateral neighbors in this order

# k off of gtp at the seem
k_offgtp_seem=[k_offgtpbase*numpy.exp(dG_latgtp), k_offgtpbase*
   numpy.exp(dG_latgtp*0.5), k_offgtpbase, k_offgtpbase*numpy.exp
   (-0.5*dG_latgtp),k_offgtpbase*numpy.exp(-dG_latgtp)]# for 0
   lateral dimers, 1/2 dimer, 1 dimer, 1.5 laterl dimers and 2
   lateral dimers

# off rates for different numbers of lateral neighbors for GDP
dG_latgdp=1.5 # lateral bond energy per bond per dimer
# ... no lateral neighbor:
k_offgdpbase=143.4 # basic rate for all dimers 1/s
k_offgdp=[k_offgdpbase*numpy.exp(dG_latgdp), k_offgdpbase,
   k_offgdpbase*numpy.exp(-dG_latgdp)]#, 702.8, 350]# 1/s; for 0
   lateral neighbors, 1 lateral neighbor
# and 2 lateral neighbors in this order

# k_off of gtp at the seem
```

```
k_offgdp_seem=[k_offgdpbase*numpy.exp(dG_latgdp), k_offgdpbase*
    numpy.exp(0.5*dG_latgdp), k_offgdpbase, k_offgdpbase*numpy.exp
    (-0.5*dG_latgdp), k_offgdpbase*numpy.exp(-dG_latgdp)]# for 0
    lateral dimers, 1/2 dimer, 1 dimer, 1.5 laterl dimers and 2
    lateral dimers
# ----------------------------------------------------------------
# reaction rates WITH VIM
# ----------------------------------------------------------------
dG_vim=2.3 # kBT energy added by bonding to a vim IF
k_offgtp_v=numpy.multiply(k_offgtp,numpy.exp(-dG_vim))
k_offgtp_seem_v=numpy.multiply(k_offgtp_seem,numpy.exp(-dG_vim))

# interaction probability of vim and IF
p_i=0.33

# rescue rate
rates_resc=1.8/60 # 1/s; only during depolymerization
rates_resc_v=9/60#1.8/60*numpy.exp(dG_vim)

# different states of the tubulin dimer have the following values:
state_gmpcpp=3
state_gdp=2
state_gtp=1
state_unoccupied=0

# number of runs of simulations
nr_simrun=100

#start timer
start = time.time()

for count_simrum in range(nr_simrun):

    # ------------------------------------------------
    # create empty microtubule
    # ------------------------------------------------
    # empty microtubules
    mt=numpy.zeros((nr_pf,max_mt_l))

    # fill first nr_seed rows with dimers in the GMPCPP state,
        this is the seed
    mt[0:nr_pf,0:nr_seed-1]=state_gmpcpp # first index is nr of
        protofilament, i.e. between 0 and nr_pf
    # second index is the number of the "layer" of tubulin dimer
        depending on the polymerization

    # set layer of gdp tubulin
    mt[0:nr_pf, nr_seed-1:nr_seed-1+lgdpshaft]=state_gtp

    # set four GTP tubulin layers at the tip
    mt[0:nr_pf, lgdpshaft + nr_seed-1:nr_seed+lgtptip-1+lgdpshaft
        ]=state_gtp

    # set start time to 0
    times=[0] # in seconds

    # set start length to 0
```

```
    l_dimers=0 # in dimer length

    # start counter of iterations
    k=0

    # initialize starting point of tip
    idx_tip=0

    # observe dimer length and check whether they are the same
    nrlsame=2000
    l_dimers_prev=numpy.zeros(nrlsame)

#    # state of microtubule:
#    depolym_check=0 # =0 : microtubule is not in rapid
   disassembly state; =1:
#    # microtubule is in rapid disassembly state
    pf_depol=1000

    #-------------------------------------------------------------
    # calculate all possible reaction rates
    #-------------------------------------------------------------
    while times[len(times)-1] < 900: # set time of produced data ;
        simulate for 600 s = 10 minutes
        # empty rates array for polymerization
        rates_polym=numpy.zeros(nr_pf)

        # empty index which dimer should poylmerize
        idx_to_pol=(numpy.zeros(nr_pf)).astype(int)

        # empty rates array fro depolymerization
        rates_depol=numpy.zeros(nr_pf)

        # save index for depolymerization / which dimer could
            depolymerize
        idx_to_dep=(numpy.zeros(nr_pf)).astype(int)

        # initalize hydrolysis indices of dimers which can be
            hydrolyzed
        idx_to_hydr=[]

       # calculate hydrolysis reaction rates:
       # find all instances where the gtp dimer is not hydrolyzed,
            but there is a GTP dimer on top
        for c in range(nr_pf): # look for two gtp dimers in a
            protofilament in a row in the GTP state
            for m in range(numpy.sum(numpy.where(mt[c,:]>0, 1,0))
                -1): # find all incorporated dimers
                if mt[c,m]==state_gtp and mt[c,m+1]>0: # test
                    whether there are two GTP dimers on top of each
                    other
                    idx_to_hydr.append((c,m)) # collect all dimers
                        which can be hydrolyzed

        # convert list of indices with dimers to hydrolyze to
            array
        idx_to_hydr=numpy.array(idx_to_hydr)

        # number of possible hydrolysis events
```

```
    nr_tohydr=idx_to_hydr.shape[0]

    # define hydrolysis rates
    rates_hydr=numpy.zeros(nr_tohydr)

    # make an array with hydrolysis rates
    for c in range(nr_tohydr): # iterate through all dimers to
        hydrolyze
        if idx_to_hydr[c, 1]<idx_tip-1:
            rates_hydr[c]=k_hy
        else:
            rates_hydr[c]=k_hy

  # calculate polymerization rate at certain spots at the end
      of MT for GTP dimer
    for c in range(nr_pf): # find first empty spot in
        protofilament which could be occupied with a new dimer
        idx_to_pol[c]=numpy.sum(numpy.where(mt[c,:]>0, 1,0)) #
            save the position where the MT polymerizes
        # check if seem
        if c==0:
            nr_neighbors=numpy.sum([(mt[nr_pf-1, idx_to_pol[c
                ]-2]>0).astype(int), (mt[nr_pf-1, idx_to_pol[c
                ]-1]>0).astype(int), (mt[1, idx_to_pol[c]]>0).
                astype(int)])
            rates_polym[c]=k_on_seem[nr_neighbors]
        if c==nr_pf-1:
            nr_neighbors=numpy.sum([(mt[0, idx_to_pol[c]+2]>0)
                .astype(int), (mt[0, idx_to_pol[c]+1]>0).astype
                (int), (mt[nr_pf-2, idx_to_pol[c]]>0).astype(
                int)])
            rates_polym[c]=k_on_seem[nr_neighbors]
        # check the number of lateral neighbor dimers at these
            positions
        if c>0 & c<nr_pf-1:
            nr_neighbors=numpy.sum([(mt[(c-1)%nr_pf,
                idx_to_pol[c]]>0).astype(int), (mt[(c+1)%nr_pf,
                 idx_to_pol[c]]>0).astype(int)])
            rates_polym[c]=k_on[nr_neighbors]

    # calculate depolymerization rate:
    for c in range(nr_pf): # find last dimer in protofilament
        which could depolymerize
        idx_to_dep[c]=(numpy.sum(numpy.where(mt[c,:]>0, 1,0))
            -1).astype(int) # this finds the first dimer
            counted from the protofilament tip
        # determine whether an IF is bound to this specific
            last dimer
        r_iv=random.random()

        # check whether this dimer is a GMPCPP dimer or not,
            if not, it is ok, it can depolymerize, if it is
            GMPCPP it will stay on bc it is the seed
        if mt[c, idx_to_dep[c]]!=state_gmpcpp:
```

```
            # check if seem
            if c==0:
                # calculate number of lateral neighbor dimers
                nr_neighbors=numpy.sum([(mt[nr_pf-1,
                    idx_to_dep[c]-2]>0).astype(int), (mt[nr_pf
                    -1, idx_to_dep[c]-1]>0).astype(int), (mt[1,
                     idx_to_dep[c]]>0).astype(int)*2])
                if mt[c, idx_to_dep[c]]==state_gtp: # if the
                    dimer is a GTP dimer
                    if r_iv<p_i: # IF and MT interact
                        rates_depol[c]=k_offgtp_seem_v[
                            nr_neighbors]
                    else:
                        rates_depol[c]=k_offgtp_seem[
                            nr_neighbors]
                if mt[c, idx_to_dep[c]]==state_gdp: # if the
                    dimer is a GDP dimer
                    rates_depol[c]=k_offgdp_seem[nr_neighbors]
            if c==nr_pf-1:
                nr_neighbors=numpy.sum([(mt[0, idx_to_dep[c
                    ]+2]>0).astype(int), (mt[0, idx_to_dep[c
                    ]+1]>0).astype(int), (mt[nr_pf-2,
                    idx_to_dep[c]]>0).astype(int)*2])
                if mt[c, idx_to_dep[c]]==state_gtp: # if the
                    dimer is a GTP dimer
                    if r_iv<p_i: # IF and MT interact
                        rates_depol[c]=k_offgtp_seem_v[
                            nr_neighbors]
                    else:
                        rates_depol[c]=k_offgtp_seem[
                            nr_neighbors]
                if mt[c, idx_to_dep[c]]==state_gdp: # if the
                    dimer is a GDP dimer
                    rates_depol[c]=k_offgdp_seem[nr_neighbors]
            if c>0 & c<nr_pf-1:
                # calculate number of lateral neighbor dimers
                nr_neighbors=numpy.sum([(mt[(c-1)%nr_pf,
                    idx_to_dep[c]]>0).astype(int), (mt[(c+1)%
                    nr_pf, idx_to_dep[c]]>0).astype(int)] )
                if mt[c, idx_to_dep[c]]==state_gtp: # if the
                    dimer is a GTP dimer
                    if r_iv<p_i: # IF and MT interact
                        rates_depol[c]=k_offgtp_v[nr_neighbors
                            ]
                    else:
                        rates_depol[c]=k_offgtp[nr_neighbors]
                if mt[c, idx_to_dep[c]]==state_gdp: # if the
                    dimer is a GDP dimer
                    rates_depol[c]=k_offgdp[nr_neighbors]

    # unit all rates / all reactions that can happen to
        perform Gillespie step
```

```
rates=numpy.concatenate((rates_hydr, rates_polym,
   rates_depol), axis=0)

# number of possible reactions
nr_reaction=rates.shape[0]

# initalize helping variables for random number selection
# random numbers for reaction time determination
R_i=numpy.zeros(nr_reaction)

# time calculated from reaction rates and random numbers
t_i=numpy.zeros(nr_reaction)

# for each element within the rates array draw a random
   number
for c in range(nr_reaction): # iterate each element
    R_i[c]= random.random()  # draw random number for each
        reaction rate
    t_i[c]=-numpy.log(R_i[c])/rates[c] # calculate time as
        in VanBurenPaper

# find index of the smallest time that passes
idx_next=numpy.argmin(t_i)

# check whether this next reaction would be addition to
   the depolymerizing protofilament
if idx_next-nr_tohydr==pf_depol and mt[pf_depol,
   idx_to_dep[pf_depol]]==state_gdp:
    t_i[idx_next]=1000 # set the reaction to an
       exaggerated time
    idx_next=numpy.argmin(t_i) # so that another reaction
       occurs instead

# check where this index is and carry out the
   corresponding step
# check whether the microtubule is in the locked
   depolymerization mode
if idx_next<nr_tohydr: #hydrolization occurs
    mt[idx_to_hydr[idx_next,0], idx_to_hydr[idx_next,1]]=
       state_gdp       # change the state of this dimer to
        GDP
else:
    if idx_next<nr_tohydr+nr_pf: #polymerization occurs
        # initialize help indices
        idx1=idx_next-nr_tohydr# number of protofilament
        idx2=idx_to_pol[idx1]# level of dimer
        mt[idx1, idx2]=state_gtp # set polymerized dimer
            to gtp state
    else: #depolymerization occurs
        if mt[idx_next-nr_tohydr-nr_pf, idx_to_dep[
           idx_next-nr_tohydr-nr_pf]]==state_gdp:# or mt[
           idx_next-nr_tohydr-nr_pf, idx_to_dep[idx_next-
           nr_tohydr-nr_pf]-1]==state_gdp:
            # only block this protofilament for growth,
                all other are allowed to grow further
```

```
                r_iv=random.random() #choose random number for
                    vim interaction
                pf_depol=idx_next-nr_tohydr-nr_pf #
                   protofilament for depolymerization
                if r_iv<p_i: # IF and MT interact
                    t_i_resc=-numpy.log(random.random())/
                       rates_resc_v
                else:
                    t_i_resc=-numpy.log(random.random())/
                       rates_resc
#                              # all depolymerization times until
    rescue
                t_i_wr=numpy.append(t_i[nr_tohydr+nr_pf:
                   nr_tohydr+2*nr_pf], t_i_resc)
                idx_next=numpy.argmin(t_i_wr)+nr_tohydr+nr_pf
                   # find fastes depolymerization rate or
                   rescue rate

            if idx_next<nr_tohydr+2*nr_pf:
                # set depolymerized dimer to unoccupied state
                mt[idx_next-nr_tohydr-nr_pf, idx_to_dep[
                   idx_next-nr_tohydr-nr_pf]]=state_unoccupied
            else: # if there is something weird in the code
                #rescue occurs
                idx_startrescue=numpy.min(numpy.where(mt[
                   pf_depol, 0:max_mt_l]==0))
                mt[pf_depol, idx_startrescue]=state_gtp
                pf_depol=1000
                #print('something went wrong')

    if l_dimers<10:
          pf_depol=1000

    # progress the reaction time
    if idx_next<len(t_i):
        times.append(times[len(times)-1]+ t_i[idx_next])
    else:
        times.append(times[len(times)-1]+ t_i_wr[len(t_i_wr)
           -1])

    if numpy.mod(k, 1000)==0:
        print(k)
        print(times[len(times)-1])

    # calculate time-grown length of MT and add the length to
       the previously saved length
    l_dimers=numpy.argmin(numpy.sum((mt[:,:]>0).astype(int),
       axis=0)>nr_pf-1)
    l_dimers_prev[nrlsame-1]=l_dimers
    l_dimers_prev[0:nrlsame-1]=l_dimers_prev[1:nrlsame]

    # check whether all elements are the same
    checklength=all(l_dimers_prev[0] == rest for rest in
       l_dimers_prev)
```

```
        if checklength and l_dimers>50:
            sys.exit('MT length stays the same')

        f=open(savename+'l_dimers_'+str(count_simrum)+'.txt','a+')
        f.write(str(l_dimers)+"\n")
        f.close()

        nr_gdps=numpy.array(numpy.where(numpy.sum((mt[:,:]==2),
            axis=0)>nr_pf-5))
        # calculate tip position
        if nr_gdps.size==0:
            idx_tip=0
        else:
            idx_tip=numpy.max(nr_gdps)

        k=k+1

    # for-loop for number of simulation steps ends

    # ---------------------------------------------------------
    # analyze catastrophe events etc
    # ---------------------------------------------------------
    # read in dimer length
    f= open(savename+'l_dimers_'+str(count_simrum)+'.txt');
    l_dimers=f.readlines()
    l_dimers=(numpy.array(l_dimers)).astype(int)


    # plot length over time
    plt.figure(count_simrum)
    plt.plot(times[0:len(l_dimers)], l_dimers[0:len(l_dimers)])
    plt.ylabel('number of dimers')
    plt.xlabel('time (s)')
    plt.savefig(savename+'time-length_'+str(count_simrum)+'.png',
        dpi=600)
    plt.close()

    # calculate catastrophe frequency
    # determine differences
    l_diff=numpy.array(l_dimers[1:len(l_dimers)])-numpy.array(
        l_dimers[0:len(l_dimers)-1])
    # find all events where there is a castrophe and sum them up
    rate_catastrophe=numpy.sum((l_diff<0).astype(int))/times[len(
        times)-1]

    # calculate growth rate
    rate_growth=numpy.sum(l_diff[l_diff>0])/times[len(times)-1]

    if count_simrum==0:
        os.mkdir(savename + 'meta_data')

    # save dimer lengths, catastrophe rate, growth rate and time
    numpy.savetxt(savename+'meta_data/time_'+str(count_simrum)+'.
        txt', times, delimiter=',')
```

```
    numpy.savetxt(savename+'rate_catastr_and_growth_'+str(
        count_simrum)+'.txt', (rate_catastrophe, rate_growth),
        delimiter=',')
    numpy.savetxt(savename+'meta_data/length_'+str(count_simrum)+'
        .txt', l_dimers, delimiter=',')

end = time.time()
time_taken = end - start
print('Time: ',time_taken)
```
