## Supplementary figures and images for "Vimentin Intermediate Filaments Stabilize Dynamic Microtubules by Direct Interactions"

### Supplementary Movie 1

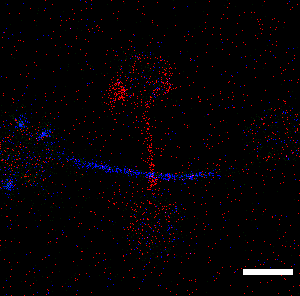

### Supplementary Movie 2

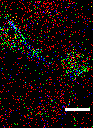

### Supplementary Movie 3

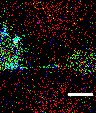

### Supplementary Movie 4

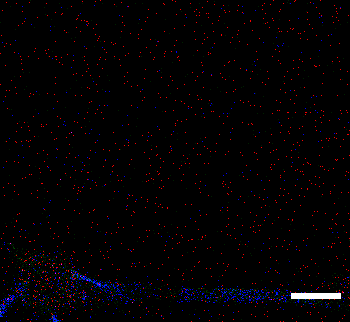

### Supplementary Movie 5

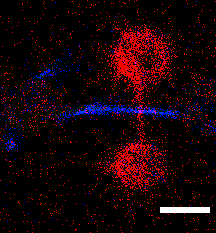

### Supplementary Movie 6

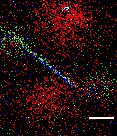
